# Asymmetry and Allostery: Insights into the Mechanism of Directional Peptide Translocation in AAA+ Unfoldases

**DOI:** 10.64898/2026.07.27.741010

**Authors:** Tyler G. Southam, Joe Breslin, Myongin Oh, Inhadul Haque, Lauren Todd, Steven E. Glynn, Jessica M. J. Swanson

**Affiliations:** Department of Chemistry, University of Utah, 315 S 1400 E, Salt Lake City, UT 84112-0850, USA; Department of Chemistry, Memorial University of Newfoundland, Core Science Facility, 45 Arctic Ave, St. John’s, NL, Canada A1C5S7; Department of Biochemistry and Cell Biology, Stony Brook University, Stony Brook, NY 11794-5215, USA

**Author notes:** Corresponding author: Jessica M. J. Swanson, **Email:**.

**Keywords:** AAA+ ATPases, Motor Protein, Unfoldase, Proteostasis, Kinetic Asymmetry, Nonequilibrium Kinetics, Mechanochemical Coupling

## Abstract

Molecular motors convert stochastic ATP chemistry into directional motion. Modern nonequilibrium theories explain this through kinetic asymmetry, yet the molecular mechanisms remain unresolved. Cryo-EM of asymmetric helical staircases has inspired the processive hand-over-hand mechanism, while biochemical and single-molecule studies have reported variable step sizes indicative of stochastic behavior. Reconciling these observations and explaining how directional motion emerges from stochastic ATP hydrolysis remain central questions. Here we use all-atom simulations and biochemical assays of AAA+ unfoldases Yme1 and Vps4 to explore structural changes induced by hydrolysis, ADP release, and ATP association. The simulations reveal pre- and post-hydrolysis conformations consistent with cryo-EM densities and interconnected allosteric communication networks. One network links the Walker B glutamate to the proximal subunit’s nucleotide binding pocket (NBP), providing a mechanism for backward propagation of hydrolysis competence around the hexamer. Another couples NBP and salt bridge rearrangements to pore-loop dynamics, connecting subunit position rather than ATP hydrolysis to substrate release. Mutational analysis demonstrates that disruption of the second network inhibits substrate degradation more than ATP hydrolysis, supporting its role in mechanochemical coupling. A second mutant points to interactions between N424 and the Walker B glutamate in ATP hydrolysis. Simulations of nucleotide exchange intermediates show that ATP binding before ADP and/or Mg^2+^ release can induce strain leading to alternative hydrolysis-competent configurations while preserving the directional bias of the staircase. Together, these findings suggest that structural asymmetry creates position-dependent allosteric interactions that generate kinetic asymmetry, biasing a fundamentally stochastic ATPase cycle toward directional substrate translocation while still permitting variations in stepping behavior.

**Significance Statement:** AAA+ ATPases play a central role in biology, converting ATP binding, hydrolysis, and release into the directional forces needed to carry out work, such as substrate translocation, remodeling, and degradation. In addition to being important therapeutic targets, understanding how these molecular machines convert stochastic ATP chemistry into persistent mechanical motion is of fundamental interest. Using simulations and biochemical assays, we identify position-responsive conformational changes and allosteric communication networks that propagate hydrolysis competence, coordinate substrate release, and redistribute the relative probabilities of mechanochemical transitions around the ATPase ring. These findings provide molecular insight into how structural asymmetry and kinetic redistribution enable directional substrate translocation.

## Introduction

Molecular motors convert stored chemical free energy into directional motion and mechanical work. Among the most powerful of these are AAA+ unfoldases, ring-shaped molecular machines capable of gripping folded proteins and generating forces sufficient to unfold, remodel, or degrade substrates as they are translocated through a central pore.^1–5^ Cryo-electron microscopy (cryo-EM) studies of multiple AAA+ unfoldases have revealed a remarkably conserved architecture in which the five or more, but most frequently six, ATPase domains adopt an asymmetric helical staircase around a bound substrate peptide (**Fig. 1**).^1, 6–10^ Four or five subunits typically engage the substrate through conserved pore-loop interactions, while the remaining subunits occupy transitional positions at the top and bottom of the staircase.

**Figure 1.**
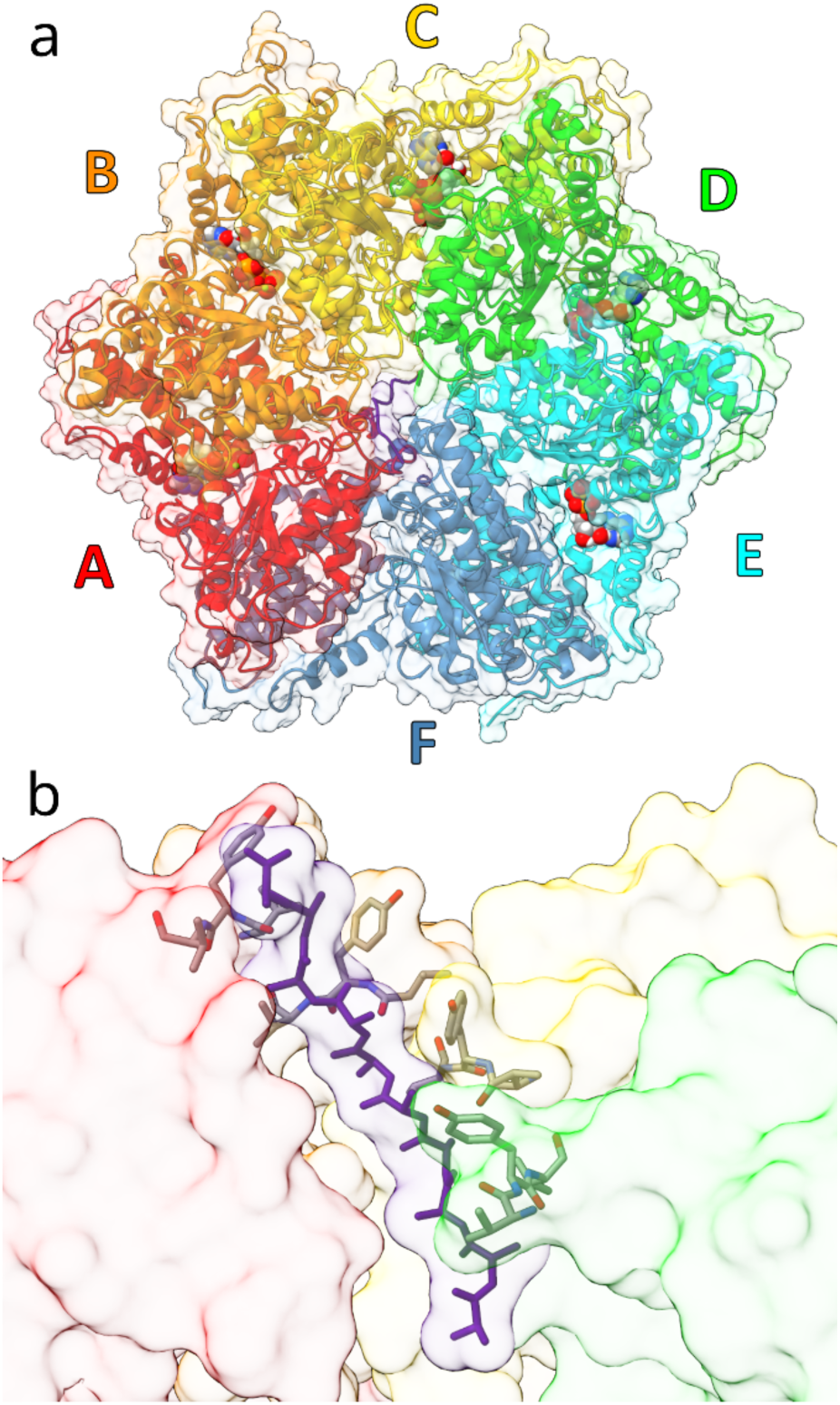
Hexameric Yme1 AAA+ unfoldase. **a)** Top-down view of Yme1 showing ATP bound in the NBPs of subunits A, B, and C; ADP bound in the NBPs of subunits D and E; and a poly-alanine substrate peptide bound in the central pore. **b)** Cutaway side-view of the central pore showing the bound substrate peptide and interacting pore loop 1 residues (V353, Y354, and V355) from subunits A through D.

These structures have inspired the processive hand-over-hand model in which ATPase subunits cycle sequentially through distinct positions around the hexamer. Each engaged subunit grips approximately two substrate residues as it descends the spiral staircase. Following ATP hydrolysis and substrate release near the bottom of the staircase, the disengaged subunit resets to the top, binds ATP, re-engages the next two substrate residues, and advances the staircase by one position, producing net substrate translocation of ∼2 amino acids per ATPase cycle.^1, 11–17^

The hand-over-hand model provides a compelling structural framework for how inter-subunit motions can produce substrate translocation. It does not, however, explain how these motions are biased to proceed in one direction consistently enough to induce forces sufficient to remodel proteins. Each ATPase subunit contains the same conserved nucleotide-binding motifs and interacts with neighboring subunits through equivalent structural elements. ATP, ADP, P_i_, Mg^2+^, and substrate binding and release are all reversible processes, and the substrate itself provides little intrinsic directional bias beyond resisting translocation. How chemically identical ATPase domains nevertheless acquire distinct mechanochemical functions that bias these reversible transitions toward forward translocation remains a fundamental unresolved question.

At the same time, biochemical and single-molecule studies have demonstrated behavior that is more complex than a strictly repeating hand-over-hand cycle. Variable step sizes, bursts of translocation, slips, pauses, bidirectional translocation and substrate-dependent behavior have been observed in multiple AAA+ systems.^18–22^ Moreover, engineered hexameric rings containing hydrolysis-incompetent subunits retain substantial translocation activity, demonstrating that productive substrate movement can occur without obligatory sequential hydrolysis by every subunit.^23, 24^ These findings have motivated models emphasizing probabilistic nucleotide turnover, rapid conformational fluctuations, and Brownian-ratchet behavior.^22, 23, 25–29^ Together, these observations suggest that although the staircase can strongly bias directional translocation, multiple microscopic mechanochemical pathways remain accessible.

Modern nonequilibrium theories have fundamentally reshaped how molecular motors are interpreted.^30–32^ Rather than invoking deterministic power strokes, these frameworks show that directional motion emerges because chemical energy biases the probabilities of microscopic transitions while preserving microscopic reversibility. In this view, kinetic asymmetry—the unequal probability of competing forward and reverse transitions within a reaction network—generates net probability current and directional motion under nonequilibrium conditions.

Together, these structural, biochemical, and theoretical advances raise a common question: how does the conserved asymmetric staircase physically construct the kinetically biased reaction network that produces directional translocation in some ATPases, and bidirectional translocation in others?^22^ More specifically, what residue-level interactions differentiate successive staircase positions, how are hydrolysis-associated conformational changes communicated between neighboring ATPase sites, and how are nucleotide-state transitions coordinated with substrate association, release, and staircase reset?

To help address these questions, we combine all-atom molecular dynamics simulations, biochemical assays, and cryoENsemble^33^ analysis of the AAA+ unfoldases Yme1^17^ and Vps4^11^ to reconstruct conformational transitions spanning one ATPase cycle. Comparison of these structurally conserved but functionally distinct unfoldases reveals how differences in inter-domain interactions, conformational stability, substrate orientation, and backbone versus side-chain interactions likely tune substrate affinity and functional specialization while preserving a common mechanochemical framework. The simulations identify previously unresolved mechanochemical intermediates and residue-level allosteric networks that couple neighboring nucleotide binding pockets (NBPs) between subunits and with substrate-binding pore loops. Mutation analyses further verify the role of these networks in directional translocation from the ATPase to the protease domain in Yme1. Together, these results provide molecular insights into the motions that link a conserved asymmetric staircase to position-dependent transition probabilities and the redistribution of kinetic asymmetry needed to drive directional substrate translocation.

## Results

### 1) Conserved staircase architecture supports different dynamics and substrate engagement in Yme1 and Vps4

Simulations were initiated from cryo-EM structures of Yme1 (PDB 6AZ0)^17^ and Vps4 (PDB 6AP1)^11^, which exhibit remarkably similar asymmetric helical staircase architectures despite serving distinct biological functions (**Fig. 1**). In both structures, ATP/Mg^2+^ is bound in subunits A-C, ADP is bound in E, and F is nucleotide free. Subunit D is bound by ADP in Vps4 and ATP/Mg^2+^ in Yme1. Subunits A–E engage a substrate peptide through conserved pore-loop interactions, while subunit F occupies a transition state between the bottom and top of the staircase. The NBP is formed at the interface between neighboring subunits and contains highly conserved Walker A (K327, T328), Walker B (E381), and arginine-finger (R435, R438) motifs (**Fig. 2**). In the simulations, multiple alternative nucleotide-bound states were run for a total of ∼40.7 μs as described in Supporting Information and summarized in **Table S1**. We varied ATP/Mg^2+^ vs. ADP in subunit D (referred to as YLT and YLD herein), ATP association to subunit F (ATP-F), Mg^2+^-retention in subunit E, and substrate orientations.

**Figure 2.**
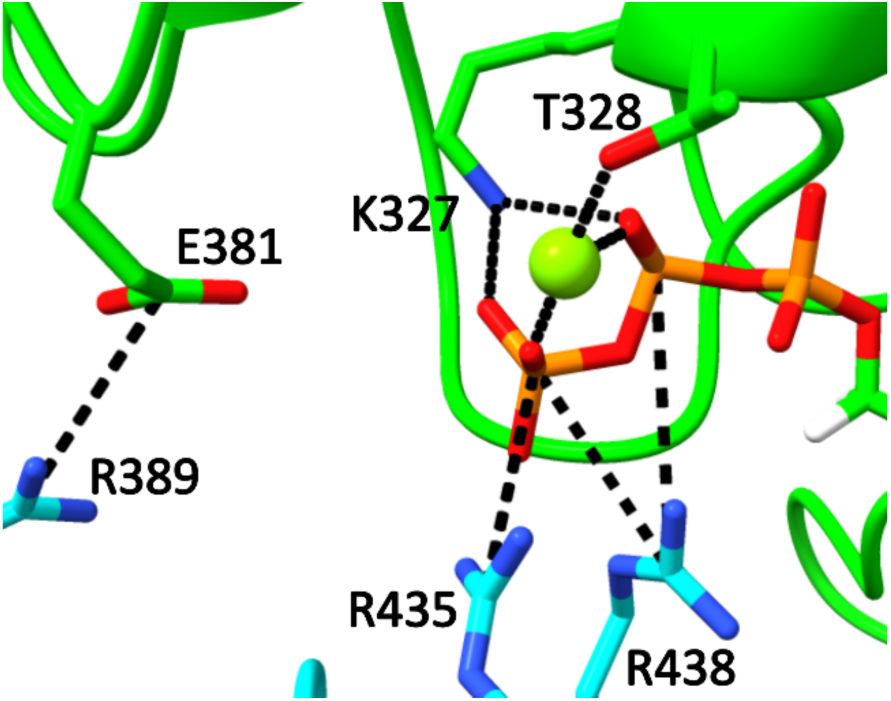
Nucleotide binding pocket of subunit D (green) with ATP bound highlighting canonical interactions, including Walker A (K327, T328) coordinating phosphate oxygens and Mg^2+^ (green sphere); Walker B (E381) in a salt-bridge with R389 from the next subunit (cyan) (note Walker B D380 is not shown for clarity); and the arginine fingers (R435, R438) from the next subunit stabilizing ATP phosphates.

We observed clear differences in the structural stability of the unfoldases: although both retained their overall staircase architecture throughout the simulations, Yme1 exhibited substantially greater stability than Vps4 (**Fig. S1**). Removal of the Yme1 protease domain increased flexibility to levels comparable to Vps4, verifying that the protease ring mechanically stabilizes the ATPase assembly. This distinction is consistent with their biological roles. Yme1 functions as a constitutive membrane protease that must process a diverse range of substrates while maintaining a stable translocation channel. Vps4 acts in ESCRT remodeling, where dynamic assembly and disassembly of the hexamer is regulated by an additional ESCRT interaction domain and cofactors to ensure ATPase activity only happens when and where it is needed.^34^ Other AAA+ ATPases that engage substrates in diverse cellular locations also co-associate with cofactors, such as ClpB and co-chaperones DnaK and DnaJ. Thus, the roles of cofactors in directional translocation is an important topic of future research.

There were also similarities and differences in how Vps4 and Yme1 engage the substrate peptide. Although structural characterization has largely focused on the stacking of enzyme and substrate side chains, we find that both unfoldases engage the substrate primarily via stable backbone-backbone hydrogen bonds (**Fig. 3**). This is consistent with their ability to translocate a wide range of sequences. However, side chain interactions can additionally modulate the stability of engagement (**Fig. S2**). Additionally, the bond patterns differ for the two unfoldases: Vps4 uses residues M205 and K207 from adjacent subunits to bind a single substrate residue, while Yme1 uses residues V353 and V355 from the same subunit to bind the same substrate residue (**Fig. 3**) while the pore loop 1 Y354 stacks between substrate side chains (**Fig. S3**). The preferred orientations are also different. Short simulations of reversed substrate orientations suggested that Vps4 preferentially engages the C-terminus of ESCRT substrates, consistent with recent substrate affinities.^34^ Yme1 more readily accommodates N-terminal engagement (**Fig. S4**), although substrate-specific degron recognition may further modulate these preferences.^5, 11, 35^

**Figure 3.**
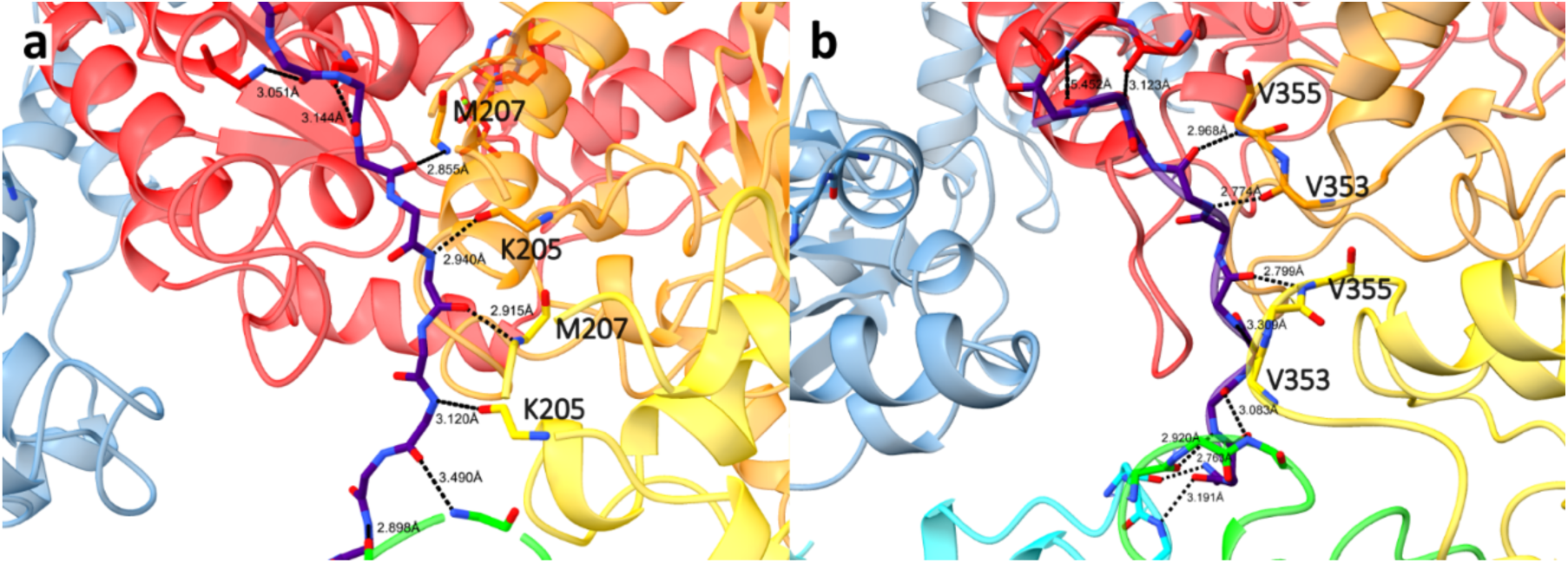
Vps4 vs Yme1 pattern of pore loop 1 hydrogen bonds with substrate. **a)** Vps4 binds the C-terminus of the substrate peptide, placing it near subunit E in the simulated structure and the N-terminus near subunit A. The NH and CO groups from the same substrate residue form hydrogen bonds with M205 and K207 from two neighboring Vps4 subunits. The backbone atoms of residues 205-207 are shown in stick style. **b)** Yme1 binds the N-terminus of the substrate peptide, resulting in the opposite orientation compared to Vps4. Additionally, the NH and CO groups from the same substrate residue form backbone hydrogen bonds with V353 and V355 from the same Yme1 subunit. The backbone atoms of residues 353-355 are shown in stick style.

Given the challenges of converging results for the more flexible Vps4, the mechanistic analyses below focus on Yme1. These simulations do not explicitly model hydrolysis or Pi/Mg^2+^ release; the possible implications of this limitation of this are considered in the Discussion. To explore if multiple nucleotide-bound states contribute to the cryo-EM structure, the simulation ensembles were first compared to the reported Yme1 density. We used the cryoENsemble^33^ method to identify the weighted combinations of specific configurations that best reproduced the experimental density map (**Fig. 4, Fig. S5**). In line with the structure containing ATP in subunit D, we find that ATP-bound (YLT) conformations contributed more strongly than ADP-bound (YLD) conformations, suggesting that both the wild-type enzyme and the hydrolysis-arrested E381Q mutant used to obtain the structure populate a larger fraction of pre-hydrolysis states. However, ADP-bound post-hydrolysis states contribute ∼35.7% while ATP-bound to subunit-F conformations contribute over 10%, consistent with ATP rebinding during reset of the staircase. Together, these observations indicate that the cryo-EM density reflects a population average over multiple mechanochemical intermediates rather than a single nucleotide-bound state.

**Figure 4.**
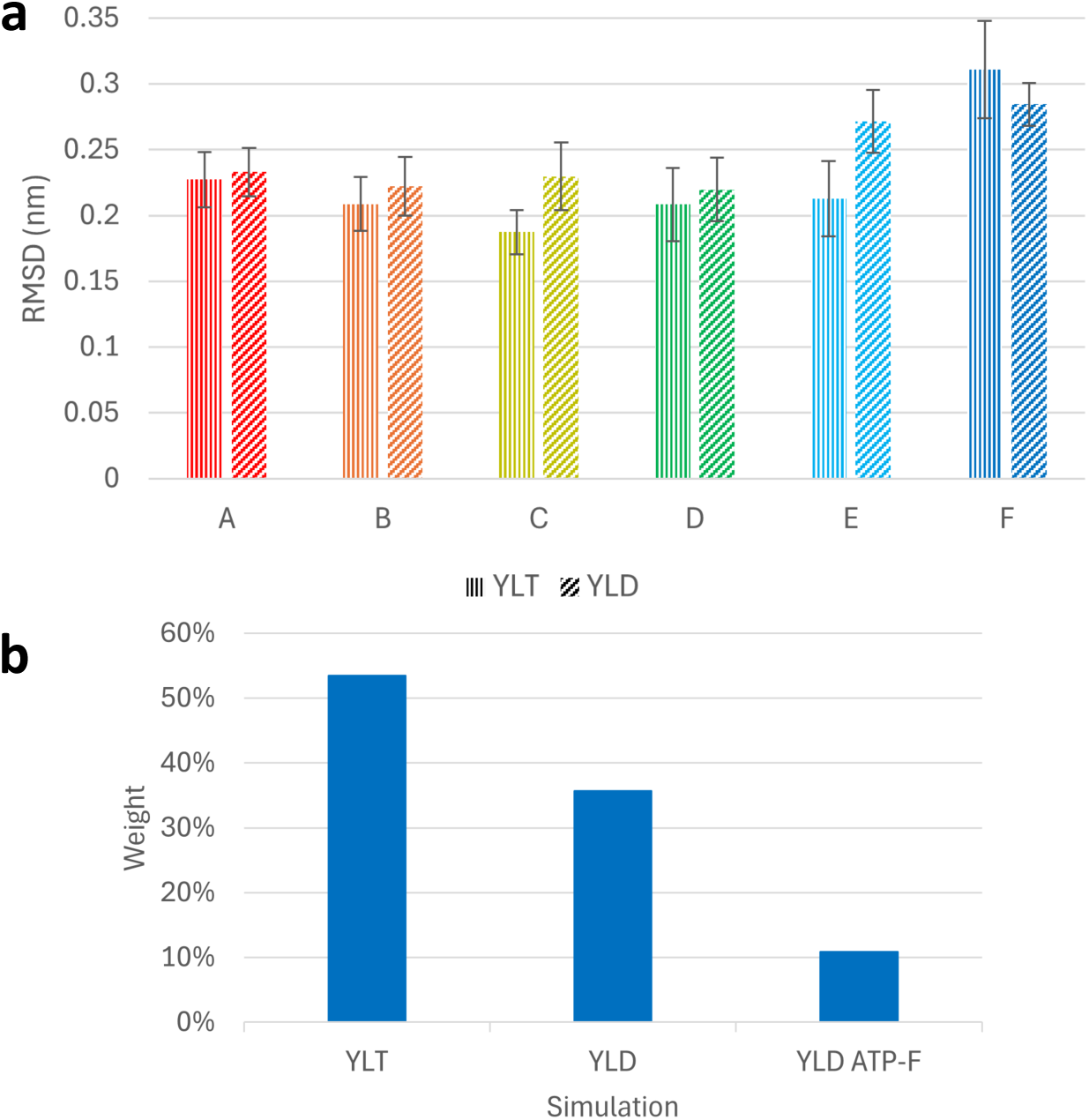
RMSD and cryoENsemble measurements. **a)** Average RMSD relative to the cryo-EM structure for each subunit in <u>Y</u>me1 with a poly<u>L</u> substrate and either A<u>T</u>P or A<u>D</u>P bound subunit D (YLT or YLD, respectively). Bars depict fluctuations measured by +/- 1 standard deviation. Values were calculated from data after 350 ns simulation time for YLT and after 400 ns simulation time for YLD, corresponding to when each system’s RMSD converged. **b)** Cumulative cryoENsemble weights of frames from each simulation that best reproduce the cryo-EM density. 21 weighted frames are represented: 11 from YLT, 7 from YLD, and 3 from YLD ATP-F.

### 2) Four position-dependent NBP conformations differentiate ATPase-cycle states

Comparison of ATP-bound and ADP-bound NBPs revealed four recurring conformational states associated with progression through the ATPase cycle (**Fig. 5**). Two pre-hydrolysis states were distinguished by the interaction between the Walker B glutamate E381 and the next subunit residue R389. In the first state (**Fig. 5a**), E381 forms a stable intersubunit salt bridge with R389 while ATP remains tightly coordinated by Walker A residues and the arginine fingers. In the second state (**Fig. 5b**), observed predominantly in subunit D, this interaction elongates, freeing E381 to adopt a hydrolysis-competent orientation. In the first post-hydrolysis state (**Fig. 5c**), primarily found in subunits D and E, release of inorganic phosphate allows E381 to move into the NBP, disrupting the E381–R389 interaction and coinciding with retraction of the arginine fingers. A second post-hydrolysis state (**Fig. 5d**) emerges in subunits E and F, where E381 has transitioned to form a previously unrecognized salt bridge with the arginine finger R435. This interaction stabilizes an open NBP compatible with ADP release and ATP rebinding. Although the new E381-R435 salt-bridge is not in the cryo-EM structure, it is present in 9 out of the 21 cryoENsemble frames that best reproduce the cryo-EM density (7 out of 7 from YLD, 1 out of 11 for YLT, and 1 out of 3 for YLD ATP-F), suggesting that it is present in the structural ensemble, but not sufficiently dominant to be reflected in structures fit to represent the ensemble as a whole. Together, these four conformational states occur during successive stages of the ATPase cycle and provide a structural framework for distinguishing key intermediates.

**Figure 5.**
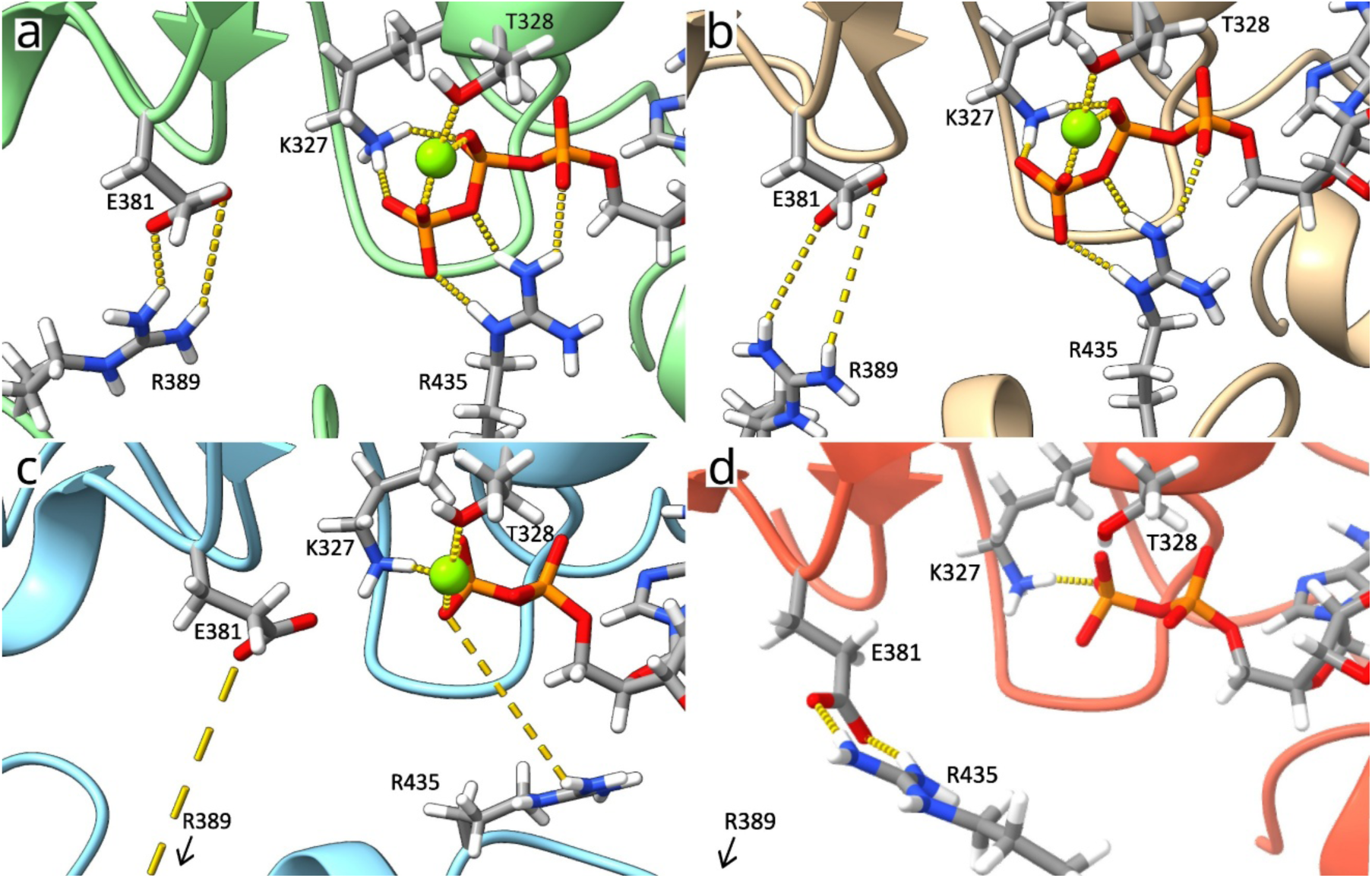
Four NBP conformations associated with successive stages of the ATPase cycle.**a)** Snapshot of YLT subunit B NBP showing the first pre-hydrolysis state characterized by the E381– R389 salt bridge and stable interactions between ATP and residues K327, R435, and R438 (R438 not shown for image clarity). **b)** Snapshot of YLT subunit D NBP showing the second pre-hydrolysis state, differentiated by the lengthening of the E381–R389 salt bridge. **c)** Snapshot of YLD subunit D NBP showing the first post-hydrolysis state. Note the complete loss of the E381– R389 salt bridge with E381 oriented towards ADP and R389 rotated out of the frame. Also note R435’s rotation away from the ADP phosphate groups. **d)** Snapshot of YLD subunit E NBP showing the second post-hydrolysis state. R389 remains rotated out of frame. R435 has formed a new salt bridge with E381.

### 3) An intersubunit allosteric network propagates hydrolysis-associated conformational changes around the staircase

A processive hand-over-hand mechanism requires ATP hydrolysis and the ensuing conformational changes to occur preferentially in specific positions within the helical staircase. Consequently, hydrolysis-associated dynamics in one NBP must influence the conformational state of neighboring ATPase sites. Having identified four recurring NBP conformations associated with successive stages of the ATPase cycle (**Fig. 5**), we next examined how transitions between these states are communicated around the hexamer.

Comparison of the pre- and post-hydrolysis simulation ensembles indicates that ATP hydrolysis is accompanied by a reorganization of interactions involving the Walker B glutamate E381. This residue starts in a stable salt bridge with R389 from the next subunit (subunits A-C); moves to a more mobile position capable of interacting with N424 (**Fig. S7**) and participating in catalysis (occasionally in subunit C but primarily in subunit D); and then moves further into the NBP post-hydrolysis to engage the arginine-finger residue formerly stabilizing the γ-phosphate of ATP (subunits E and F) (**Fig. 5a-c**). This movement of E381 also displaces a loop containing R389 in the same subunit, which is salt-bridged to E381 from the previous subunit (**Fig. S6)**. Deformation of the R389 loop weakens the E381-R389 interaction in the preceding subunit. Thus, hydrolysis in one ATPase site mechanically perturbs the neighboring NBP through a shared intersubunit interaction network.

Although the preceding subunit consistently experienced weakening of the E381-R389 interaction, we did not always observe the large-scale separation of E381 from R389. For example, following hydrolysis in subunit D, subunit C generally retained the E381-R389 interaction, although it shifts between pre-hydrolysis states 1 and state 2 (**Fig. 5a,b**). In contrast, the lower staircase positions (D to E transition) underwent substantially larger rearrangements that disrupted and reoriented the E381-R389 interaction network (**Fig. 5c**). The final transition to the second post-hydrolysis state (**Fig. 5c**) involves further reorganization of the E381-R389 network with E381 forming a new salt bridge with R435 (**Fig. 5d**). This interaction appears to stabilize a late-stage post-hydrolysis configuration in transitioning subunits E and F while simultaneously removing E381 from participation in the E381-R389 interaction network.

This allows the network to transmit smaller, potentially hydrolysis-enabling conformational changes to the preceding NBP, while complete reorganization of the interface is limited to progression through the lower staircase. Together, these observations identify an intersubunit allosteric network centered on E381 and R389 that transmits hydrolysis-associated rearrangements backward around the hexamer.

### 4) Lower-staircase rearrangements couple NBP opening to substrate release

Communication between neighboring NBPs does not, by itself, explain how unfoldase cycling produces directional substrate translocation. In particular, substrate release must be associated with the bottom of the staircase to maintain a directional force, even if hydrolysis occurs in higher subunits. We next examined how the larger conformational changes in the lower subunits influence the substrate-binding pore loops.

The position-dependent responses described above coincide with distinct conformations of the R389 loop, which is extended in the substrate-engaged subunits A–D, begins to reorganize in E, and substantially shortened in F as pore loop 2 folds into the N-terminus of helix α5. Closer analysis revealed a second allosteric communication pathway extending from R389 to pore loop 2 (**Fig. 6**). R389 forms persistent interactions with N402 and Q399 at the base of pore loop 2 and immediately adjacent to the conserved substrate-contacting pore loop 2 residue Y396. Thus, R389 lies at the intersection of two communication pathways: one linking neighboring NBPs and a second linking the NBP to the substrate-binding interface.

**Figure 6.**
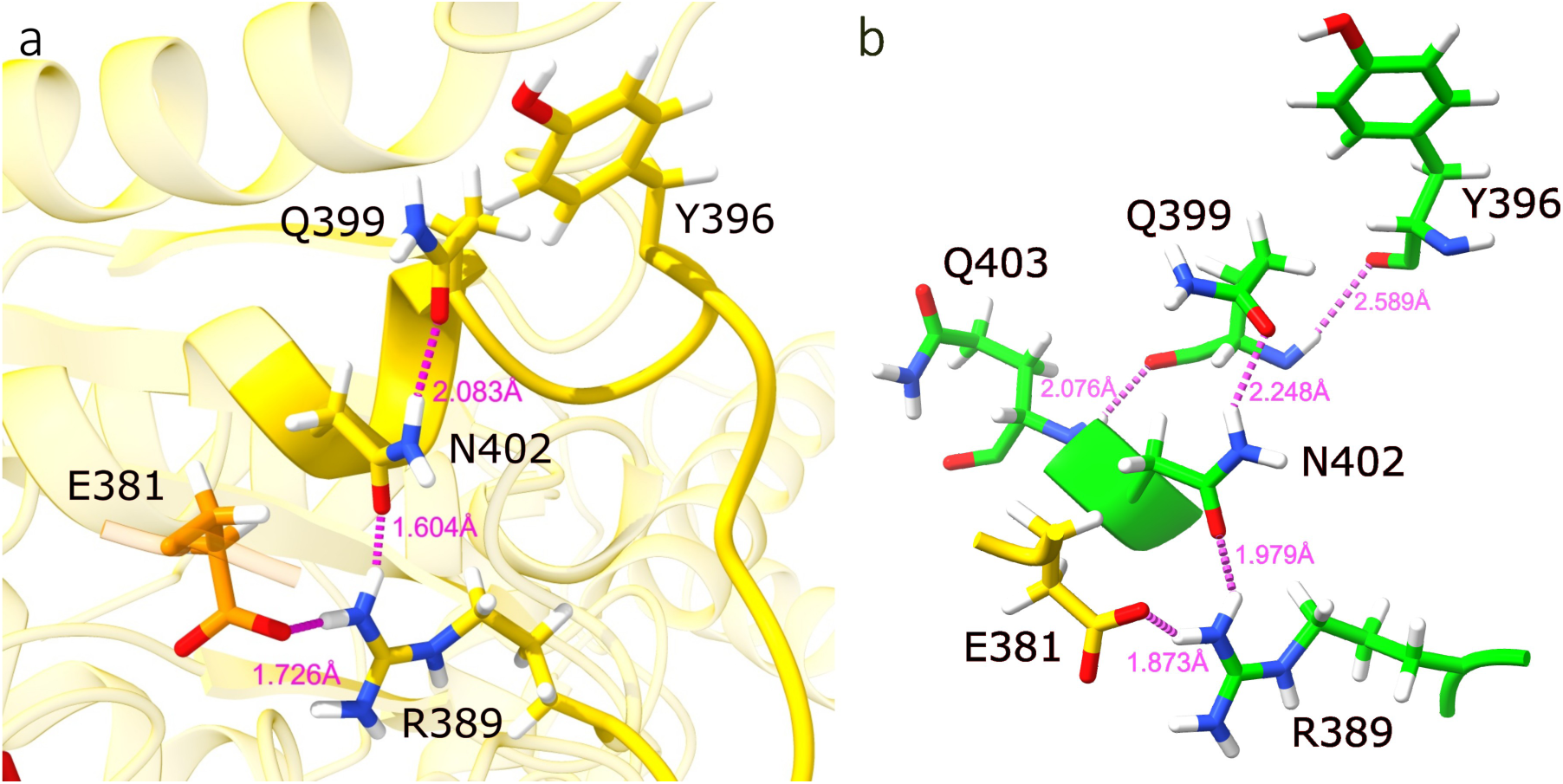
Hydrogen bond network linking the NBP to the substrate peptide. **a)** The primary network from the Walker B E381 residue, through R389, N402, and Q399. The nearby pore loop 2 Y396 residue is shown to illustrate its proximity. **b)** The redundant backbone hydrogen bonds network incorporating Q403.

Throughout most of the staircase, substrate engagement was maintained by backbone-mediated hydrogen bonds between pore loops and the substrate peptide (**Fig. 3**) complemented by Y354 stacking between substrate side chains (**Fig. S3**). In contrast, the lowest staircase positions, particularly subunit E, displayed increased fluctuations in substrate contacts and more probable release for handoff to the protease. Similar pore-loop geometries were observed across polyalanine, polyleucine, and mixed-sequence substrates despite differences in interaction energies and residence times. Thus, substrate sequence can modulate the strength and lifetime of engagement, while release is preferentially coupled to progression into the lowest staircase positions.

Together, these observations show that substrate release is primarily governed by staircase position and by the large conformational rearrangements of the lowest subunit. Opening of the lowest NBP can weaken substrate contacts through the R389–Q399–N402–pore-loop interface, among other interactions, while allowing substrate engagement to persist if hydrolysis occurs earlier or higher in the ring.

### 5) Experimental verification of allosteric network

To test the proposed allosteric network linking the NBP to pore-loop 2, three substitutions were introduced into ^cchex^Yme1, a soluble, active enzyme construct, and assayed for effects on both ATP hydrolysis and substrate degradation.^36^ Initial degradation rates were determined from proteolysis of unfolded, fluorescein-labeled Tim10, a known Yme1 substrate (Fig. 7).^35^

**Figure 7.**
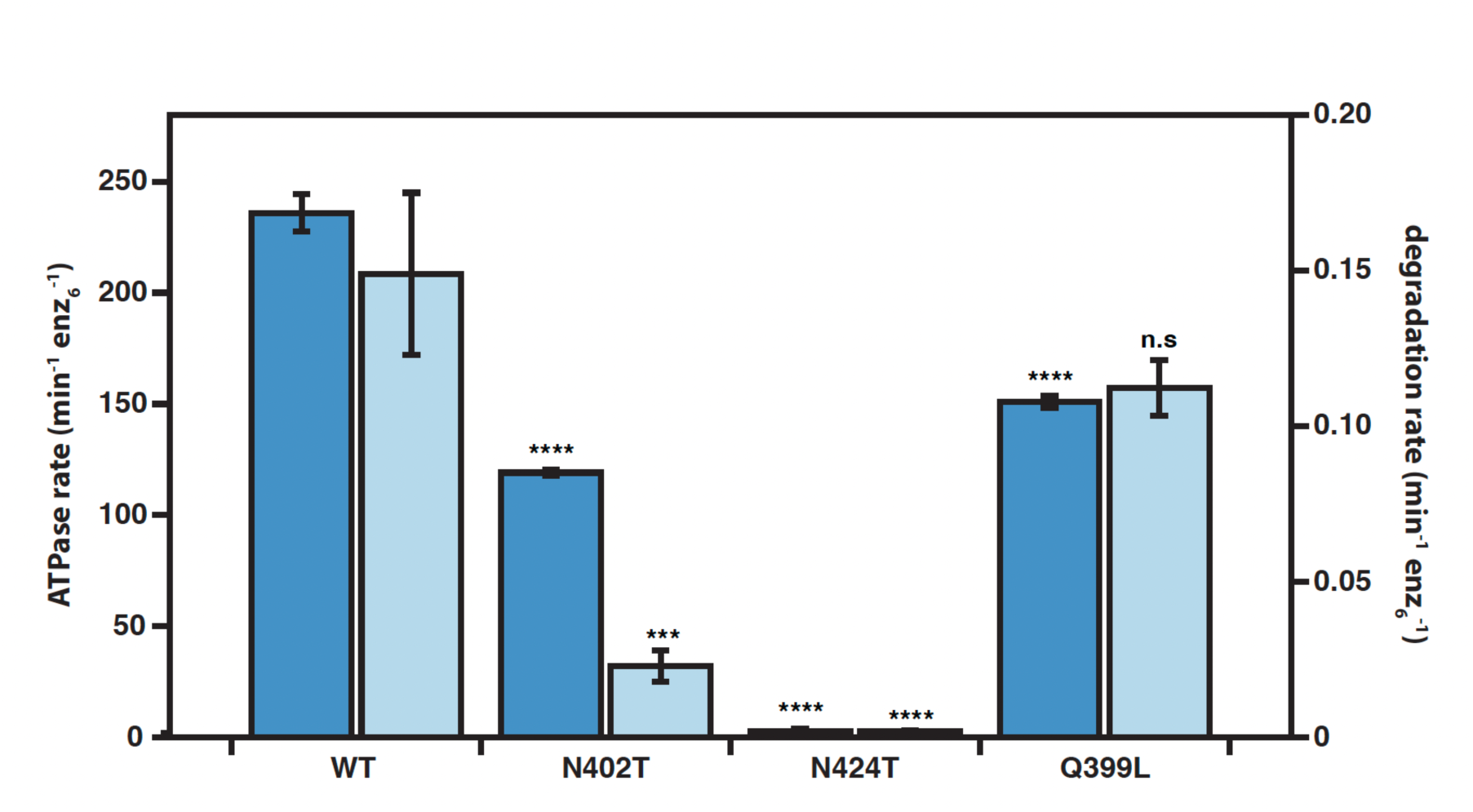
ATPase and protein degradation rates for ^cchex^Yme1 and its variants validate the functional relevance of the allosteric network between NBP and pore-loop 2. Values are means of independent replicates (n ≥ 3) ± SD. ***P ≤ 0.001, ****P ≤ 0.0001 as calculated using Student’s two-tailed t test and statistical significances are shown relative to WT ^cchex^Yme1.

N402T was chosen because this residue links the motion of R389 to Q399 through two hydrogen bonds; the shorter Thr side chain of the mutant can satisfy only one of these interactions at a time and is predicted to interrupt the pathway. N402T retained ∼50 % of wild-type ATPase activity but only ∼15 % of the wild-type degradation rate (Fig. 7). This disproportionate loss of degradation is consistent with ATP hydrolysis becoming uncoupled from translocation of substrate into the protease chamber. Notably, N402 corresponds to N432 of human AFG3L2, where the equivalent N432T substitution, which displays a similar reduction in ATPase and degradation activity, is linked to the development of spinocerebellar ataxia type 28.^1, 37^ The loss of coupling between ATP hydrolysis and degradation in both enzymes suggests that the allosteric network is shared among AAA+ proteases.

As an alternative disruption of the same network Q399L was tested, but this variant retained ∼60 % ATPase and ∼75 % degradation activity. Closer inspection reveals additional hydrogen bonds linking backbone atoms of Q399 with both Q402 and Y396, indicating redundancy within this network that can preserve partial degradation in the mutant. Lastly, N424T was selected because this residue is highly conserved and lies close to E381 in the second pre-hydrolysis and first post-hydrolysis states, suggesting a role in positioning E381 for catalysis following its release from R389. Indeed, interactions of E381 with N424 and R389 are anticorrelated in subunit D. (**Fig. S7**) Before hydrolysis, E381 alternates coordination between R389 and N424. Following hydrolysis, the E381-R389 salt bridge is fully lost while N424 stably engages E381. In agreement with a key role in catalysis, N424T completely abolished both ATP hydrolysis and degradation activity (Fig. 7).

#### Staircase position also controls nucleotide binding and release

Another key to directional translocation in AAA+ ATPases is ATP preferentially binding at the top of the staircase and ADP/Mg^2+^/Pi release at the bottom. For this to occur, there must be position-dependent factors differentiating association affinities within the NBPs. Consistent with this, ADP was released from the NBP in subunit E, following hydrolysis in D and the ensuing conformational changes. Release occurred after 1.3 μs in the YLD simulation and only 1.0 μs in the ATP-F simulation.

In the upper staircase ATP-bound subunits (A-C), many canonical intersubunit interactions remain intact, including the intersubunit-signaling (ISS) motif (D409–G410–F411) (**Fig. S6**), previously proposed to sense the nucleotide state via D409 and relay it to the pore loops, initiating substrate release.^17, 38^ In addition, upper subunits maintain the Walker A–nucleotide– arginine finger (R435) coordination, and the E381–R389 salt bridge (**Fig. 5a, Fig. S8a**). In the subunit D NBP pre-hydrolysis state (**Fig. 5b, Fig. S8b**), the ISS and nucleotide interactions are maintained while the E381–R389 salt bridge fluctuates in and out of contact.

Post-hydrolysis in subunit D (**Fig. 5c, Fig. S8c**), these intersubunit interactions are disrupted: the ISS motif begins to disconnect, the arginine fingers pull away from the nucleotide, and the salt bridge permanently dissociates, with R389 eventually flipping directions to point away from subunit D and towards subunit F. While the arginine fingers dissociate from the nucleotide, they are not outside of the range of fluctuating back into contact. These changes are more pronounced in subunits E and F, but in different ways. In subunit E, the NBP has opened so far that the nucleotide can no longer bind both Walker A (K327, T328) and arginine finger residues (R435, R438) simultaneously. This facilitates ADP release in YLD and ATP-F. Opening of the NBP also pulls the ISS motif further apart (**Fig. S8**).

In subunit F, ISS residue F411 further folds into the C-terminus of helix α5 (**Fig. S8d**). R389 is still flipped to point away from the NBP towards subunit A. As subunit F packs onto the top of the hexameric ring, the Walker A and arginine finger residues are again close enough to simultaneously bind ATP, partially due to the salt bridge between E381 and R435, while the ISS motif and E381–R389 salt bridge are close enough to begin interacting again (**Fig. S8e**).

These distinct residue rearrangements make it possible for ATP to stably bind in the subunit F NBP and for the F-to-A interface to reform, stabilizing intersubunit interactions. This is not the case in subunit E, where key intersubunit interactions are lost (e.g., F411 from the ISS is stably wound up in helix α5). Additionally, any bound nucleotide can only contact either the Walker A motif or the arginine fingers, but not both, reducing its stability in the pocket. This leads to a preferential binding of ATP to subunit F and preferential loss of ADP from subunit E. The presence of ADP in subunit E would also reduce the opportunity for ATP to bind to the lowest subunit and pull the hexameric ring backward. These observations provide a molecular explanation for how the conformational rearrangements in the lowest and resetting subunits result in preferential association of ATP at the top of the hexameric staircase and release of ADP from the bottom.

### 6) Perturbed nucleotide ordering produces strained but translocation-competent intermediates

The allosteric networks described above operate within a structurally asymmetric staircase in which each subunit occupies a distinct conformational environment. A global measurement of these differences is apparent in the ATPase-protease angle previously identified by Puchades et al.^17^ as a descriptor of staircase geometry. Consistent with the cryo-EM structures, simulations revealed a progressive increase in ATPase-protease angle from subunit A through subunit E, followed by a sharp transition in subunit F (**Fig. 8**). As the substrate moves toward the protease domain, the ATPase domains tilt progressively downward (the angle increases). The largest angles occur in subunit E, where substrate release begins, whereas subunit F adopts a distinct upward orientation consistent with reset of the staircase. This global measure is used to analyze the impact of adding ATP to subunit F and of retaining Mg²⁺ with ADP in subunit E (both with ADP bound in subunit D).

**Figure 8.**
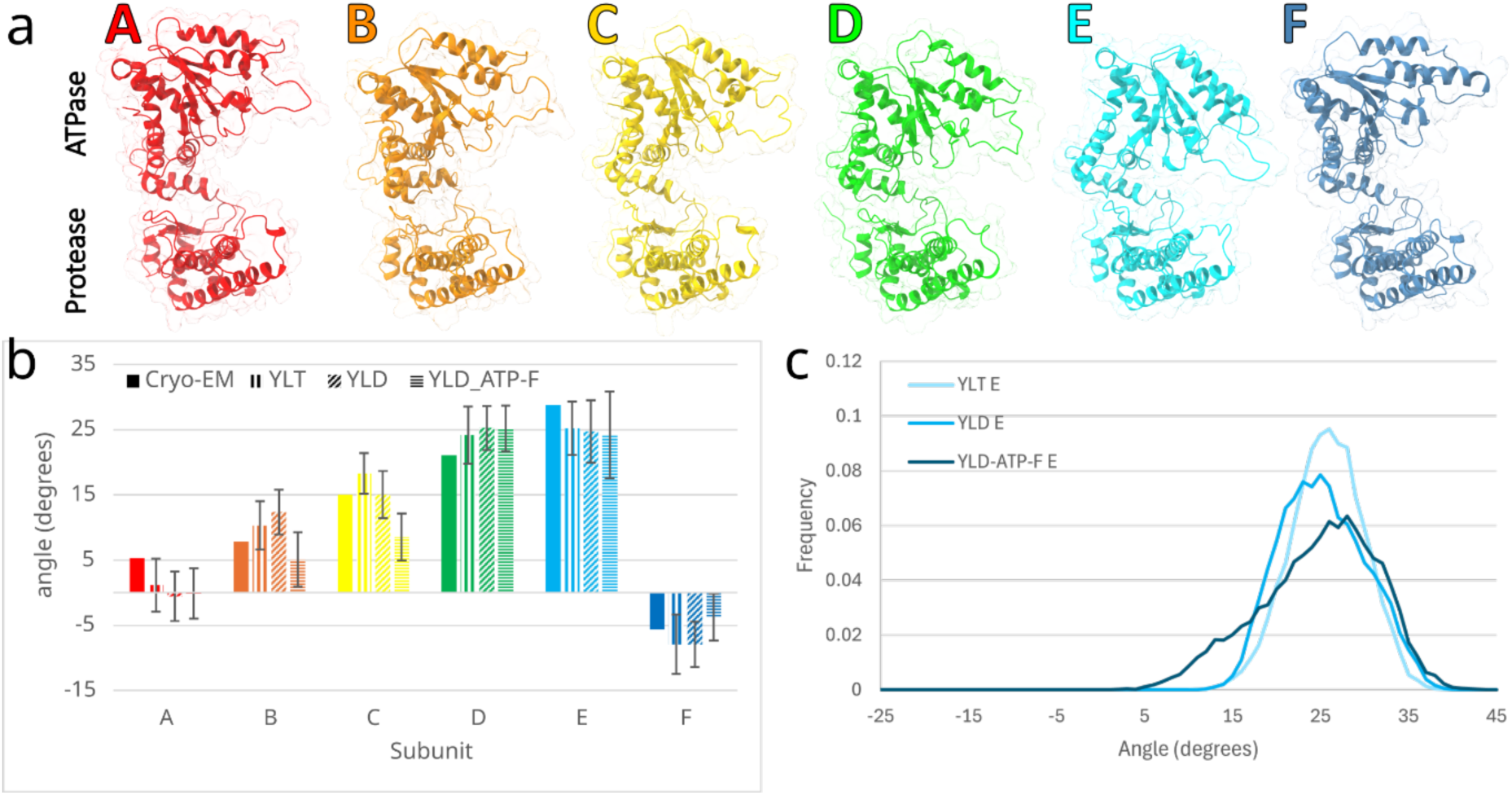
ATPase-protease angle in Yme1. Larger angles indicate a steeper downward angle of the ATPase domain towards the protease domain. **a)** Representative views of subunits A through F from the YLT simulation aligned by protease domains. **b)** The angle from the initial Cryo-EM structure and average angles from the YLT, YLD, and YLD ATP-F simulations. Error bars are estimated errors calculated with autocorrelation and block averaging using GROMACS’ analyze function. **c)** Angle distributions for subunit E in the YLT, YLD, and YLD ATP-F simulation trajectories.

First, analyzing the ATP-F simulations, we find that following the addition of ATP, subunit F forms preliminary contacts with the substrate above subunit A, while subunit E loses persistent substrate interactions (**Figs. S3, S9**). Simultaneously, ATP binding promotes closure of the F NBP and restoration of interactions characteristic of upper staircase subunits. These observations are consistent with a hand-over-hand cycle in which ATP binding re-engages the resetting subunit at the top of the spiral as the lowest engaged subunit releases the substrate.

The simulations further suggest that structural asymmetry constrains the order of nucleotide turnover. In the YLT and YLD simulations, the ATPase-protease angles followed a relatively smooth progression around the ring (**Fig. 8b**). In contrast, ATP binding to subunit F before complete turnover of subunit E generated distortions in this progression and increased conformational heterogeneity. Similar effects were observed in trajectories retaining Mg²⁺ within the E NBP after hydrolysis. Because Mg²⁺ stabilizes nucleotide coordination, its retention delays relaxation of the post-hydrolysis state and alters the conformational transitions required for reset, suggesting Mg²⁺ is released with P_i_, consistent with the resolved structure.

The consequences of this perturbation extend beyond individual ATPase sites. ATP-F simulations exhibited loss of intersubunit-signaling (ISS) motif interactions in ATP-bound subunits that would normally remain engaged (**Fig. S10**). Notably, disruption of ISS contacts occurred without substrate release, indicating that ISS rearrangement alone is insufficient to trigger pore-loop disengagement. Instead, ATP-F simulations appear to populate strained intermediate states in which the staircase architecture remains intact but communication between neighboring ATPase sites becomes partially frustrated.

Together, these observations suggest that the asymmetric staircase favors a preferred sequence of events. ATP hydrolysis in subunit D is followed by product release and conformational relaxation in subunit E before ATP binding fully stabilizes subunit F. Premature ATP binding or delayed product release does not destroy the staircase but instead generates additional strain within the interconnected allosteric network. Thus, structural asymmetry not only establishes distinct functional roles for different subunits but also organizes the temporal ordering (kinetic asymmetry) of hydrolysis, product release, substrate disengagement, and ATP rebinding around the ring.

### 7) An integrated mechanochemical model links nucleotide cycling to directional translocation

The conformational states and allosteric communication pathways identified above can be combined into a mechanistic proposal for one complete ATPase cycle (**Fig. 9**). Several movies illustrate this mechanism from different angles: a composite view (**Movie S1**), a side view showing the pore loop 1 motifs (**Movie S2**), a view from above the hexamer (**Movie S3**), and individual views from subunits D, E, and F (**Movies S4-S9**). This mechanistic model is necessarily provisional and will require refinement as the mechanisms of phosphate and Mg²⁺ release, as well as the contribution of the hydrolysis reaction itself, become better understood. Although individual NBPs occupy distinct conformational states at any given time, the simulations suggest a preferred progression linking ATP hydrolysis, substrate release, nucleotide exchange, and staircase reset. The model explains how structural asymmetry biases ATP hydrolysis to happen in subunit D, substrate release in subunit E, and ATP association to the F/A interface.

**Figure 9.**
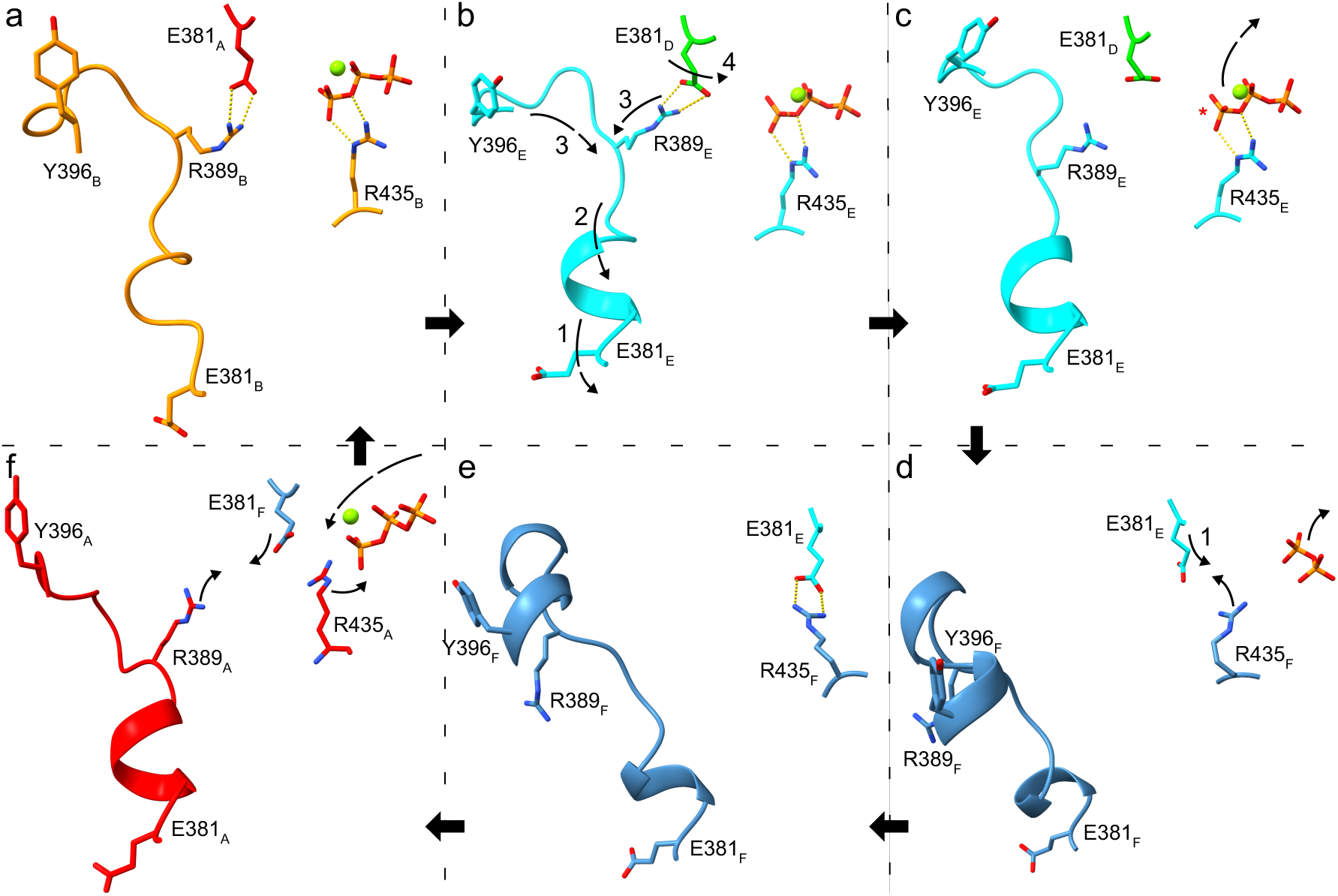
Illustration of the proposed mechanochemical cycle in Yme1. **a)** The cycle starts with the upper-staircase subunits in the stable ATP-bound pre-hydrolysis state, as shown in the subunit A NBP between subunit A (red) and subunit B (orange). This conformation will most often be carried through to subunit D’s NBP (as shown in b).**b)** Next, the inward motion of E381_E_ in subunit E (cyan) pulls R389_E_, destabilizing the salt bridge to E381_D_ in subunit D (lime) while simultaneously pulling pore loop 2. The allosteric effect of E381_E_’s motion is illustrated by arrows. **c)** Destabilization of the E381_D_-R389_E_ salt bridge then frees E381_D_ to participate in ATP hydrolysis (*), followed by the escape of P_i_ and Mg^2+^. **d)** Next, E381_E_ associates with R435_F_ and ADP escapes, as shown in the subunit E NBP between subunit E and subunit F (blue). Note that the E381-R435 salt bridge may already form in subunit D’s NBP post-hydrolysis. **e)** The E381_E_-R435_F_ salt bridge remains in the apo state. **f)** At the top of the array, ATP rebinds subunit F, reforming the E381_F_-R389_A_ and R435_A_-ATP salt bridges to subunit A (red), thus resetting the hexamer.

The cycle begins in the upper subunits with a stable E381-R389 interfacial interaction and ATP coordinated by Walker A residues and the arginine fingers (**Fig. 9a**). As subunits descend the staircase, rearrangement of the E381-R389 network in subunit E begins to elongate the E381-R389 interaction in subunit D, leading to the first and then the second pre-hydrolysis states, with the latter freeing E381 to interact with N424 and participate in catalysis (**Fig. 9b**). ATP hydrolysis and phosphate release initiate opening of the NBP in subunit D (**Fig. 9c, d**). These changes reorganize the E381-R389 interaction network, displacing the R389 loop. Because R389 forms the junction between the two allosteric pathways identified above, this rearrangement simultaneously weakens the E381-R389 interaction in the preceding subunit (C in this case) while altering the R389-Q399-N402-pore-loop network that stabilizes substrate binding. The result is propagation of hydrolysis-associated conformational changes toward the preceding ATPase site while substrate engagement is weakened only as the hydrolyzed subunit progresses into the lowest subunit position in the staircase (D to E).

Further opening of the NBP in subunit E accompanies product release and formation of the second post-hydrolysis state, in which E381 forms a stable interaction with R435 (**Fig. 9e**). This state is associated with the most open NBP, progressive loss of substrate contacts, and preparation for nucleotide exchange. Following ADP release, the apo subunit restacks onto the top of the hexamer where ATP binding stabilizes reformation of the upper staircase interactions (**Fig. 9f**). ATP binding promotes closure of the NBP while simultaneously establishing preliminary substrate contacts above subunit A, preparing subunit F for reinsertion at the top of the staircase.

## Discussion

Cryo-EM structures have transformed our understanding of AAA+ unfoldases by revealing a conserved asymmetric helical staircase architecture and inspiring the processive hand-over-hand model of substrate translocation. Our findings largely support this framework while identifying conformational intermediates and allosteric pathways that connect nucleotide turnover to conformational changes in neighboring ATPase sites and substrate-binding pore loops. The central result is that staircase position places chemically identical ATPase domains in distinct interaction environments, thereby favoring hydrolysis, product release, substrate disengagement, nucleotide exchange, and reset at different positions around the ring.

The proposed mechanism links the structural asymmetry observed in AAA+ unfoldases to the nonequilibrium principles that govern directional motion. Nonequilibrium theory establishes that reversible chemical transformations produce net motion through kinetically asymmetric reaction networks; our results identify position-dependent structural interactions that contribute to those asymmetric transition probabilities. By coupling substrate association to ATP association at the top of the helix and release to the larger motions only present at the bottom, the staircase does not require a perfectly deterministic firing sequence. Rather, it favors a dominant mechanochemical pathway with hydrolysis in subunit D, while retaining forward translocation in less probable alternatives. For example, premature ATP association or delayed product release can produce strain across the staircase that could result in hydrolysis in higher subunits while forward translocation is retained.

The comparison of Yme1 and Vps4 further illustrates how a conserved translocation architecture can be adapted to different functions. Stabilization of Yme1 by its protease ring contrasts with the greater dynamics of the regulated Vps4 assembly. In both systems, backbone-mediated substrate interactions support broad sequence tolerance, whereas side-chain contacts and preferred peptide orientation can tune affinity and substrate selectivity. Functional diversification can therefore arise through changes in assembly stability and substrate recognition without altering the underlying mechanochemical organization.

The structural intermediates identified here provide a starting framework for developing quantitative nonequilibrium kinetic models of AAA+ unfoldases, rather than a complete mechanistic description. Realizing such models will first require a more complete understanding of the chemical mechanism of ATP hydrolysis, proton transfer, and the timing and mechanochemical consequences of Pi and Mg²⁺ release through QM/MM simulations, enhanced-sampling approaches, and additional complementary experiments. Beyond these molecular details, the models will require transition rates that depend explicitly on thermodynamic driving forces, mechanical state, substrate interactions, and nucleotide occupancy to determine how these factors redistribute flux through competing mechanochemical pathways. Nevertheless, it is noteworthy that simulations of distinct nucleotide-bound states alone reproduce nearly the complete sequence of conformational changes associated with subunit reset. This finding suggests that ATP hydrolysis acts primarily by altering the molecular interactions that differentiate nucleotide and substrate association from dissociation, while the resulting position-dependent remodeling of allosteric networks, including those identified herein, drives much of the subsequent mechanical cycle.

## Conclusions

By comparing simulations of distinct nucleotide-bound intermediates, we reconstruct much of the conformational progression spanning one AAA+ unfoldase cycle. The simulations identify four recurring NBP intermediates and two interconnected allosteric communication pathways centered on R389. One pathway links hydrolysis-associated rearrangements to the preceding NBP, providing a structural mechanism for backward propagation of hydrolysis competence that biases ATP hydrolysis toward subunit D. The second couples NBP rearrangements to pore loop 2 dynamics, coordinating substrate release with progression into the lowest subunit position. Mutations of the second network validate its role in substrate release from the lowest subunit, enabling handoff to the protease domain.

Together, these findings suggest how a conserved asymmetric helical staircase can organize chemically identical ATPase domains into position-dependent interaction environments that favor different mechanochemical transitions around the ring. Rather than requiring a perfectly deterministic hydrolysis sequence, this organization accommodates stochastic excursions, including ATP hydrolysis higher in the staircase, while preserving forward progression by coupling substrate release to the larger conformational transitions of the lowest subunit and substrate association to the top of the staircase.

Comparison of Yme1 and Vps4 further shows how this conserved mechanochemical architecture can be adapted to distinct biological functions through differences in conformational stability, substrate orientation, and substrate interactions, while preserving the underlying translocation mechanism and broad substrate tolerance. Although additional work is needed to resolve ATP hydrolysis, P_i_/Mg²⁺ release, and the condition-dependent kinetics governing these transitions, the structural intermediates and allosteric pathways identified herein provide a foundation for the development of nonequilibrium models capable of quantitatively relating molecular interactions to experimentally measured substrate translocation through the framework of kinetic asymmetry.

## Methods

### MD simulations

Starting structures for Vps4 and Yme1 were obtained from RCSB PDB^39^ (PDB: 6AP1 and 6AZ0, respectively). The ADP*BeFx ATP mimic in Vps4 was converted to ATP. The missing Vps4 loop residues 365-368 were added using MODELLER.^40, 41^ Initial structures were prepared with either ADP or ATP in subunit D for each unfoldase. Additional initial structures were prepared by replacing the bound substrate with poly-leucine and poly-valine-lysine substrates using MODELLER to adjust the side chains. Orientation of the substrates (C to N vs N to C) was validated with short simulations of structures with the cryo-EM modeled substrate reversed using MODELLER.

Molecular dynamics (MD) simulations were performed with the GROMACS simulation package,^42, 43^ the ANTON2 software,^44^ and the CHARMM36m force field.^45, 46^ ^44^ANTON2 simulations were started from structures retrieved from the final frame of their respective GROMACS MD simulations, with specific modifications: an ATP·Mg^2+^ complex was placed in subunit F’s NBP, the substrate peptide was lengthened by four residues on the C terminus using MODELLER, the ADP molecule in subunit E was reinserted into the NBP in those structures where it had escaped during the GROMACS simulation and a replica with Mg^2+^ added to subunit E’s NBP was also prepared.

### Cryo-EM ensemble identification using cryoENsemble

To evaluate which nucleotide-bound states from our simulations best reflect the cryo-EM ensemble density, we used cryoENsemble,^33^ a tool developed to extract and weight structures from simulations. An initial set of 2000 representative structures from the YLT, YLD, YLD ATP-F, and YLD Mg-E simulations was obtained and fit to the cryo-EM density map. As suggested in its original implementation, cryoENsemble was run in iterative mode in two separate runs; first, weighting against a half-resolution cryo-EM density map, then against the full-resolution map. This reduced the number of simulation structures from 2000 to just 21 that best reproduced the cryo-EM density.

### Cloning, Expression and Purification

Variants of ^cchex^Yme1 were produced by site-directed mutagenesis using a previously generated 2G-T plasmid (Addgene #29707) containing the ^cchex^Yme1 open reading frame as a template.^35^ All ^cchex^Yme1 variants were expressed and purified as previously described.^5^

### Biochemical Assays

ATPase activity measurements were performed using a coupled-enzyme assay monitoring the loss of NADH absorbance at 340 nm, as previously described with some modifications.^35^ Reactions containing 0.5 µM ^cchex^Yme1 or its variants were carried out at 30 °C in a 384-well clear bottom plate (Corning) using a SpectraMax iD5 plate reader (Molecular Devices). Reactions were performed in PD buffer (25 mM HEPES-KOH (pH 7.5), 100 mM KCl, 5 mM MgCl_2_, 5% glycerol) supplemented with 1 mM NADH, 21.5 U ml^−1^ lactate dehydrogenase, 2 mM ATP, 7.5 mM phosphoenolpyruvate and 18.8 U ml^−1^ pyruvate kinase.

Degradation rates of fluorescein-labeled Tim10 (Tim10-FL) were measured by monitoring fluorescence increases (ex = 495 nm; em = 535 nm) accompanying relief of dye self-quenching associated with proteolysis. Labeled substrate was produced by incubating yeast Tim10 (∼120 µM) at 30 °C in the presence of 6 M guanidine hydrochloride and 1 mM DTT to unfold the protein and fully reduce its internal cysteine residues for labeling. DTT was removed by buffer exchange using Bio-spin desalting columns (BioRad) followed by addition of 80-fold molar excess fluorescein-5-maleimide (Sigma Aldrich) at 30 °C for 120 min. Following completion of the reaction, excess dye was removed by buffer exchange into PD buffer and substrate concentration was quantified by comparing SDS-PAGE band intensity against a standard curve of known Tim10 concentrations. Degradation reactions containing 0.5 µM ^cchex^Yme1 or its variants and 1 µM Tim10-FL in the presence of an ATP regeneration system (16 mM creatine phosphate, 0.32 mg/mL creatine kinase, and 4 mM ATP) were performed in PD buffer at 30 °C in 384-well black plates (Corning) using a SpectraMax iD3 plate reader (Molecular Devices). Initial time points with linear increases in fluorescence were used to calculate degradation rates.

### Visualization and other analyses

VMD^45^ and UCSF ChimeraX^46, 47^ were used for trajectory visualizations. GROMACS native analysis functions, our own scripts (some using the MDAnalysis^48, 49^ library), and cryoENsemble were employed for trajectory analysis. Data are presented using DataGraph.^47^

Further details of simulation parameters, systems simulated, equilibration protocols, and analyses are available in the **SI Appendix**, including **Tables S1-S4**.

## Acknowledgments

The authors thank Professor Chris Hill for many helpful discussions and for inspiring their work. They thank Dudu Tong for early work on the project. The work was supported by NIH NIGMS (R35GM143117 to J.M.J.S and R01GM115898 to S.E.G.) and by a collaborative development award from the CHEETAH Center at the University of Utah, United States of America (NIH P50 AI150464). Computational resources were provided by Expanse at the San Diego Supercomputing Center and STAMPEDE2 at the Texas Advanced Computing Center through the Advanced Cyberinfrastructure Coordination Ecosystem: Services and Support (ACCESS) program (allocation MCB200018) supported by NSF (grant nos. 2138259, 2138286, 2138307, 2137603, and 2138296); by the Anton 2 supercomputer at the Pittsburgh Supercomputing Center (PSC) through Grant R01GM116961 from the National Institutes of Health (generously made available by D.E. Shaw Research); and by the Center for High-Performance Computing (CHPC) at the University of Utah.

Molecular graphics and analyses performed with UCSF ChimeraX, which was developed with support from National Institutes of Health R01-GM129325 and the Office of Cyber Infrastructure and Computational Biology, National Institute of Allergy and Infectious Diseases.

## Author Contributions

T.S. and J.M.J.S. designed the research. T.S. performed the simulations. I.H. and L.T. performed the biochemical studies. T.S., M.O., S.E.G., and J.M.J.S. analyzed the results and wrote the manuscript.

## Competing Interest Statement

The authors acknowledge they have no competing interests.

## Notes

### Competing Interest Statement

The authors have declared no competing interest.

### Summary of Updates

This version of the manuscript includes experimental validation of the allosteric networks predicted by simulations and suggests an important role for N424 in hydrolysis.

## References

1. C. Puchades, C. R. Sandate, G. C. Lander, The molecular principles governing the activity and functional diversity of AAA+ proteins. Nat. Rev. Mol. Cell Biol. 21, 43– 58 (2019).

2. J. Snider, G. Thibault, W. A. Houry, The AAA+ superfamily of functionally diverse proteins. Genome Biol. 9, 216 (2008).

3. Y. A. Khan, K. I. White, A. T. Brunger, The AAA+ superfamily: a review of the structural and mechanistic principles of these molecular machines. Crit. Rev. Biochem. Mol. 57, 156–187 (2021).

4. T. Ogura, A. J. Wilkinson, AAA+ superfamily ATPases: common structure–diverse function. Genes to Cells 6, 575–597 (2001).

5. M. Quispe-Carbajal, L. Todd, S. E. Glynn, Recognition of small Tim chaperones by the mitochondrial Yme1 protease. Protein Sci. 35 (2026).

6. R. Strack, Cryo-EM goes atomic. Nat. Methods 17, 1175–1175 (2020).

7. S. E. Glynn, J. R. Kardon, O. Mueller-Cajar, C. Cho, AAA+ proteins: converging mechanisms, diverging functions. Nat. Struct. Mol. Biol. 27, 515–518 (2020).

8. S. N. Gates, A. Martin, Stairway to translocation: AAA+ motor structures reveal the mechanisms of ATP-dependent substrate translocation. Protein Science 29, 407–419 (2020).

9. X. Fei, T. A. Bell, S. Jenni, B. M. Stinson, T. A. Baker, S. C. Harrison, R. T. Sauer, Structures of the ATP-fueled ClpXP proteolytic machine bound to protein substrate. Elife 9 (2020).

10. S. N. Gates, A. L. Yokom, J. Lin, M. E. Jackrel, A. N. Rizo, N. M. Kendsersky, C. E. Buell, E. A. Sweeny, K. L. Mack, E. Chuang, et al., Ratchet-like polypeptide translocation mechanism of the AAA+ disaggregase Hsp104.. 357, 273–279 (2017).

11. H. Han, N. Monroe, W. I. Sundquist, P. S. Shen, C. P. Hill, The AAA ATPase Vps4 binds ESCRT-III substrates through a repeating array of dipeptide-binding pockets. Elife 6, 1–15 (2017).

12. H. Han, C. P. Hill, Structure and mechanism of the ESCRT pathway AAA+ ATPase Vps4. Biochem. Soc. Trans. 47, 37–45 (2019).

13. E. C. Twomey, Z. Ji, T. E. Wales, N. O. Bodnar, S. B. Ficarro, J. A. Marto, J. R. Engen, T. A. Rapoport, Substrate processing by the Cdc48 ATPase complex is initiated by ubiquitin unfolding.. 365 (2019).

14. I. Cooney, H. Han, M. G. Stewart, R. H. Carson, D. T. Hansen, J. H. Iwasa, J. C. Price, C. P. Hill, P. S. Shen, Structure of the Cdc48 segregase in the act of unfolding an authentic substrate.. 365, 502–505 (2019).

15. M. Su, E. Z. Guo, X. Ding, Y. Li, J. T. Tarrasch, C. L. Brooks, Z. Xu, G. Skiniotis, Mechanism of Vps4 hexamer function revealed by cryo-EM. Sci. Adv. 3 (2017).

16. S. Sun, L. Li, F. Yang, X. Wang, F. Fan, M. Yang, C. Chen, X. Li, H. W. Wang, S. F. Sui, Cryo-EM structures of the ATP-bound Vps4E233Q hexamer and its complex with Vta1 at near-atomic resolution. Nat. Commun. 8, 1–13 (2017).

17. C. Puchades, A. J. Rampello, M. Shin, C. J. Giuliano, R. L. Wiseman, S. E. Glynn, G. C. Lander, Structure of the mitochondrial inner membrane AAA+ protease YME1 gives insight into substrate processing.. 358, eaao0464 (2017).

18. J. C. Cordova, A. O. Olivares, Y. Shin, B. M. Stinson, S. Calmat, K. R. Schmitz, M. E. Aubin-Tam, T. A. Baker, M. J. Lang, R. T. Sauer, Stochastic but highly coordinated protein unfolding and translocation by the ClpXP proteolytic machine. Cell 158, 647–658 (2014).

19. M. E. Aubin-Tam, A. O. Olivares, R. T. Sauer, T. A. Baker, M. J. Lang, Single-Molecule Protein Unfolding and Translocation by an ATP-Fueled Proteolytic Machine. Cell 145, 257–267 (2011).

20. A. O. Olivares, H. C. Kotamarthi, B. J. Stein, R. T. Sauer, T. A. Baker, Effect of directional pulling on mechanical protein degradation by ATP-dependent proteolytic machines. Proceedings of the National Academy of Sciences 114, E6306–E6313 (2017).

21. 21. M. Sen, R. A. Maillard, K. Nyquist, P. Rodriguez-Aliaga, S. Pressé, A. Martin, C. Bustamante, The ClpXP protease unfolds substrates using a constant rate of pulling but different gears. Cell 155, 636 (2013).

22. R. Casier, D. Levy, I. Riven, Y. Barak, G. Haran, A stochastic mechanism drives fast substrate translocation in the AAA+ machine ClpB. Nature Communications 17, 1773- (2026).

23. A. Martin, T. A. Baker, R. T. Sauer, Rebuilt AAA + motors reveal operating principles for ATP-fuelled machines. Nature 437, 1115–1120 (2005).

24. V. Baytshtok, J. Chen, S. E. Glynn, A. R. Nager, R. A. Grant, T. A. Baker, R. T. Sauer, Covalently linked HslU hexamers support a probabilistic mechanism that links ATP hydrolysis to protein unfolding and translocation. Journal of Biological Chemistry 292, 5695–5704 (2017).

25. D. Gai, R. Zhao, D. Li, C. V. Finkielstein, X. S. Chen, Mechanisms of Conformational Change for a Replicative Hexameric Helicase of SV40 Large Tumor Antigen. Cell 119, 47–60 (2004).

26. R. Russell, A. Matouschek, Chance, Destiny, and the Inner Workings of ClpXP. Cell 158, 479–480 (2014).

27. H. Mazal, M. Iljina, I. Riven, G. Haran, Ultrafast pore-loop dynamics in a AAA+ machine point to a Brownian-ratchet mechanism for protein translocation. Sci. Adv. 7 (2021).

28. I. Riven, H. Mazal, M. Iljina, G. Haran, Fast dynamics shape the function of the AAA+ machine ClpB:Lessons from single-molecule FRET spectroscopy. FEBS J. (2022). 10.1111/FEBS.16539.

29. G. Haran, I. Riven, Perspective: How Fast Dynamics Affect Slow Function in Protein Machines. J. Phys. Chem. B 127, 4687–4693 (2023).

30. E. Penocchio, G. Gu, A. Albaugh, T. R. Gingrich, Power Strokes in Molecular Motors: Predictive, Irrelevant, or Somewhere in Between? J. Am. Chem. Soc. 147, 1063–1073 (2025).

31. E. Penocchio, G. Ragazzon, Kinetic Barrier Diagrams to Visualize and Engineer Molecular Nonequilibrium Systems. Small 19, 2206188 (2023).

32. R. D. Astumian, Kinetic asymmetry allows macromolecular catalysts to drive an information ratchet. Nat. Commun. 10, 1–14 (2019).

33. T. Włodarski, J. O. Streit, A. Mitropoulou, L. D. Cabrita, M. Vendruscolo, J. Christodoulou, Bayesian reweighting of biomolecular structural ensembles using heterogeneous cryo-EM maps with the cryoENsemble method. Sci. Rep. 14, 18149 (2024).

34. H. Wienkers, H. Han, F. Whitby, C. P. Hill, Substrate binding and coupled mechanisms of Vps4p substrate recruitment and release from autoinhibition. Sci. Rep. 15, 25024 (2025).

35. A. J. Rampello, S. E. Glynn, Identification of a Degradation Signal Sequence within Substrates of the Mitochondrial i-AAA Protease. J. Mol. Biol. 429, 873–885 (2017).

36. H. Shi, A. J. Rampello, S. E. Glynn, Engineered AAA+ proteases reveal principles of proteolysis at the mitochondrial inner membrane. Nat. Commun. 7, 13301- (2016).

37. C. Puchades, B. Ding, A. Song, R. L. Wiseman, G. C. Lander, S. E. Glynn, Unique Structural Features of the Mitochondrial AAA+ Protease AFG3L2 Reveal the Molecular Basis for Activity in Health and Disease. Mol. Cell 75, 1073–1085.e6 (2019).

38. S. Augustin, F. Gerdes, S. Lee, F. T. F. Tsai, T. Langer, T. Tatsuta, An Intersubunit Signaling Network Coordinates ATP Hydrolysis by m-AAA Proteases. Mol. Cell 35, 574–585 (2009).

39. H. M. Berman, J. Westbrook, Z. Feng, G. Gilliland, T. N. Bhat, H. Weissig, I. N. Shindyalov, P. E. Bourne, The Protein Data Bank. Nucleic Acids Res. 28, 235–242 (2000).

40. B. Webb, A. Sali, Comparative Protein Structure Modeling Using MODELLER. Curr. Protoc. Bioinformatics 54, 5.6.1–5.6.37 (2016).

41. A. Fiser, R. Kinh, G. Do, A. S. Ali, Modeling of loops in protein structures. Protein Science 9, 1753–1773 (2000).

42. M. J. Abraham, T. Murtola, R. Schulz, S. Páll, J. C. Smith, B. Hess, E. Lindah, GROMACS: High performance molecular simulations through multi-level parallelism from laptops to supercomputers. SoftwareX 1-2, 19–25 (2015).

43. H. J. C. Berendsen, D. van der Spoel, R. van Drunen, GROMACS: A message-passing parallel molecular dynamics implementation. Comput. Phys. Commun. 91, 43–56 (1995).

44. D. E. Shaw, J. P. Grossman, J. A. Bank, B. Batson, J. A. Butts, J. C. Chao, M. M. Deneroff, R. O. Dror, A. Even, C. H. Fenton, et al., Anton 2: Raising the Bar for Performance and Programmability in a Special-Purpose Molecular Dynamics Supercomputer. International Conference for High Performance Computing, Networking, Storage and Analysis, SC 2015-January, 41–53 (2014).

45. J. Huang, S. Rauscher, G. Nawrocki, T. Ran, M. Feig, B. L. De Groot, H. Grubmüller, A. D. MacKerell, CHARMM36m: an improved force field for folded and intrinsically disordered proteins. Nat. Methods 14, 71–73 (2016).

46. J. Huang, A. D. Mackerell, CHARMM36 all-atom additive protein force field: Validation based on comparison to NMR data. J. Comput. Chem. 34, 2135–2145 (2013).

47. 47. DataGraph. [Preprint] (2020). Available at: https://www.visualdatatools.com/ [Accessed 3 September 2026].

