## Supporting Information for "Asymmetry and Allostery: Insights into the Mechanism of Directional Peptide Translocation in AAA+ Unfoldases"

**This PDF file includes:**

Supporting text
Figures S1 to S11
Tables S1 to S4
Legends for Movies S1 to S9
SI References

**Other supporting materials for this manuscript include the following:**

Movies S1 to S9

### Supporting Information Text

### Detailed Methods

**Modeling of initial structures.** Starting structures were obtained from RCSB PDB[1] for Vps4 binding ESCRT-III peptide (PDB: 6AP1) and Yme1 binding a polypeptide chain (PDB: 6AZ0). In the Vps4 structure, the resolved VSL helices of the Vta1 cofactor were included; the ADP*BeFx ATP mimic was converted to ATP by replacing the BeFx group with a $PO_3$ group; and missing loop residues 365-368 were added using MODELLER.[2, 3] Residue protonation states were selected with PROPKA.[4] In Yme1, the E381Q mutation was reverted to the wild type by replacing the NE atom with an OE atom in all but one simulation that was run to compare wild type to mutant stability.

Each unfoldase is homohexameric with the subunits referred to alphabetically A-F. In both structures, subunits A to C bind ATP and a $Mg^{2+}$ ion, subunit E binds ADP, and subunit F is empty. The nucleotide bound in subunit D differed. Vps4 bound ADP, while Yme1 bound ATP and $Mg^{2+}$. This difference is consistent with the proposed hand-over-hand mechanism in which hydrolysis occurs in subunit D. Consequently, we prepared initial structures with subunit D binding ATP/ $Mg^{2+}$ or ADP for both unfoldases to examine the impact of pre- and post-hydrolysis nucleotide states on the hexamer's conformation and dynamics.

As different substrate peptide sequences have experimentally shown varying binding affinities and translocation efficiencies in various unfoldases,[5–7] additional substrate peptide sequences were modeled. In addition to the cryo-EM reported substrates, poly-leucine (polyL) and poly-valine-lysine (polyVK) -bound structures were prepared. This was accomplished using MODELLER to generate side chain coordinates from backbone positions. A two-step steepest-descent energy minimization was performed by first freezing the unfoldase and substrate backbones while allowing the side chains to optimize and then unfreezing the substrate backbone to allow the full substrate to relax.

Due to the relatively low resolution of the substrate peptides in the cryo-EM structure, the orientations (C to N or N to C) of the substrates were the result of best fitting. To validate these orientations, replicas with reversed substrates were prepared. MODELLER was again used to make this change by retaining the Cα and Cβ positions, renumbering the substrate residues in reverse order, and then building and optimizing the side chains as described above.

**Details of MD simulations.** We performed MD simulations using the GROMACS simulation package,[8, 9] the ANTON2 software,[10] and the CHARMM36m force field.[11, 12] The various simulations performed are listed in **Table S1** (including GROMACS versions used). Structures used in GROMACS MD simulations were placed in a dodecahedral box with a solvation separation of 1.5 nm between images. Structures were neutralized and solvated in 150 mM NaCl solution with the CHARMM-modified TIP3P water model.[13] The system was first minimized via steepest descent energy minimization, followed by careful multi-step equilibration to retain critical NPB interactions and give sufficient time for proper hydration. This protocol (detailed in **Tables S2-S3**) was refined after many rounds of finding that some subunits, particularly subunit B, would quickly coordinate the Walker B E381 to $Mg^{2+}$ due to lack of hydration to stabilize the divalent cation. First, an initial 100 ps "freeze" step was run, restricting all atoms except for solvent and counter-ions to allow full hydration of the $Mg^{2+}$ ions in the NBPs. Heavy-atom restraints were implemented for the next seven equilibration steps, reducing the force constant with each step from 1000 to 0 kJ/mol/nm$^2$ (**Table S2**). The first of these seven steps also used the following restraints in each applicable NBP to avoid rapid Mg2+ coordination: 1) harmonic restraints were implemented between $Mg^{2+}$ and the nucleotide phosphate groups and between $Mg^{2+}$ and T328; 2) a piecewise linear/harmonic restraint was also implemented between $Mg^{2+}$ and E381 to maintain a minimal distance of 0.45 nm (**Table S3**). Following this, equilibration steps were run using a Berendsen thermostat and/or Berendsen barostat[14] as listed in **Table S2**, with all covalent bonds to hydrogen converted to LINCS constraints.[15] GROMACS production runs were performed with an integration timestep of 2 fs using an NPT ensemble at 300 K and 1 atm with a velocity rescaling thermostat, Parrinello-Rahman barostat,[16] and with all bonds to hydrogen converted to LINCS constraints.

Structures used on the ANTON2 system were retrieved from the final frame of their respective GROMACS MD simulations. An ATP nucleotide and a $Mg^{2+}$ ion were placed in subunit F's NBP, using subunit A's NBP structure for alignment. The substrate peptide was lengthened by four residues using MODELLER by replicating the four N-terminal residues of the existing substrate and linking them to the C-terminus of the substrate. For the polyL-bound Yme1 structure, the ADP molecule was placed back in subunit E's NBP as it had escaped in the previous GROMACS production run. A replica of this system was also prepared, with the addition of a $Mg^{2+}$ ion to subunit E's NBP. After these structure modifications, a two-step vacuum optimization was run as previously described to remove substrate clashes. The structures were placed in a cubic box with a minimum buffer distance of 0.15 nm and solvated with the CHARMM-modified TIP3P water model. The system was neutralized with 150 mM NaCl and minimized via a steepest descent energy minimization. The system was equilibrated in GROMACS using the previously described freeze step, and multi-step equilibration with the addition of several restraints after the freeze step: a piecewise linear/harmonic restraint between the Cα's of the substrate peptide residues 9 and 14 was implemented to keep the substrate unfolded by maintaining a minimum distance between residues 9 and 14. Two harmonic restraints were implemented between the ATP nucleotide and K327 in the subunit F NBP. (**Table S3**) Equilibration steps used the same ensembles listed in **Table S2**.

The ANTON2 production runs were performed with an integration timestep of 2.4 fs using an NPT ensemble with a Nose-Hoover[17, 18] thermostat and Martyna, Tobias, and Klein (MTK) barostat.[19] The two harmonic restraints between ATP and K327 in subunit F were kept during the production run. These two harmonic restraints were adjusted to k ≈ 125 kJ/mol/nm for two of the systems when they crashed sometime after 7.5 μs. VMD[20] and UCFS ChimeraX[21, 22] were used for trajectory visualizations. GROMACS native analysis functions and our own scripts, some using the MDAnalysis[23, 24] library, were employed for trajectory analysis.

**Ensemble identification with cryoENsemble.** To evaluate which nucleotide-bound states from our simulations best reflect the cryo-EM ensemble density, we used cryoENsemble,[25] a tool developed to extract and weight structures from simulations. First, we identified representative structures using k-means clustering, resolving 500 clusters from the YLT, YLD, YLD ATP-F, and YLD Mg-E simulations. The initial frame of each simulation was fit to the cryo-EM density map obtained from RCSB PDB using ChimeraX's "fit in map" tool. The four 500-frame trajectories were then aligned on the Cα atoms to these map-fit initial frames. As suggested in its original implementation, cryoENsemble was run in iterative mode, weighting our 2000 cluster frames first to a half-resolution cryo-EM density map prepared with ChimeraX. This resulted in the 2000 structures being pared down to 55 weighted frames for a second iterative weighting to the full-resolution cryo-EM density map. After the second iterative run, 21 weighted frames remained. A recommended contour level of 0.08 and the resolution of the map (3.40 Å) were obtained from the validation report in the PDB entry.

**ATPase-protease angle.** Consistent with Puchades et al.,[26] the ATPase-protease angle for each subunit was defined as the angle of declination of the ATPase domain α4 and α5 helices from the plane of the protease domain (**Fig. S11**). An ATPase domain vector was calculated from the average of two vectors formed by the α4 and α5 helices. The individual helix vectors were defined by two α carbons on the same side of the helix near each end. The protease domain normal vector was the average vector of six vectors defined by the α13 helices in the protease domain. These helix vectors were also defined by pairs of α carbons. The angle between each ATPase domain vector and the protease normal vector was then calculated. Subtracting from 90° yields an angle of declination from the protease normal vector's plane.

**Figures**

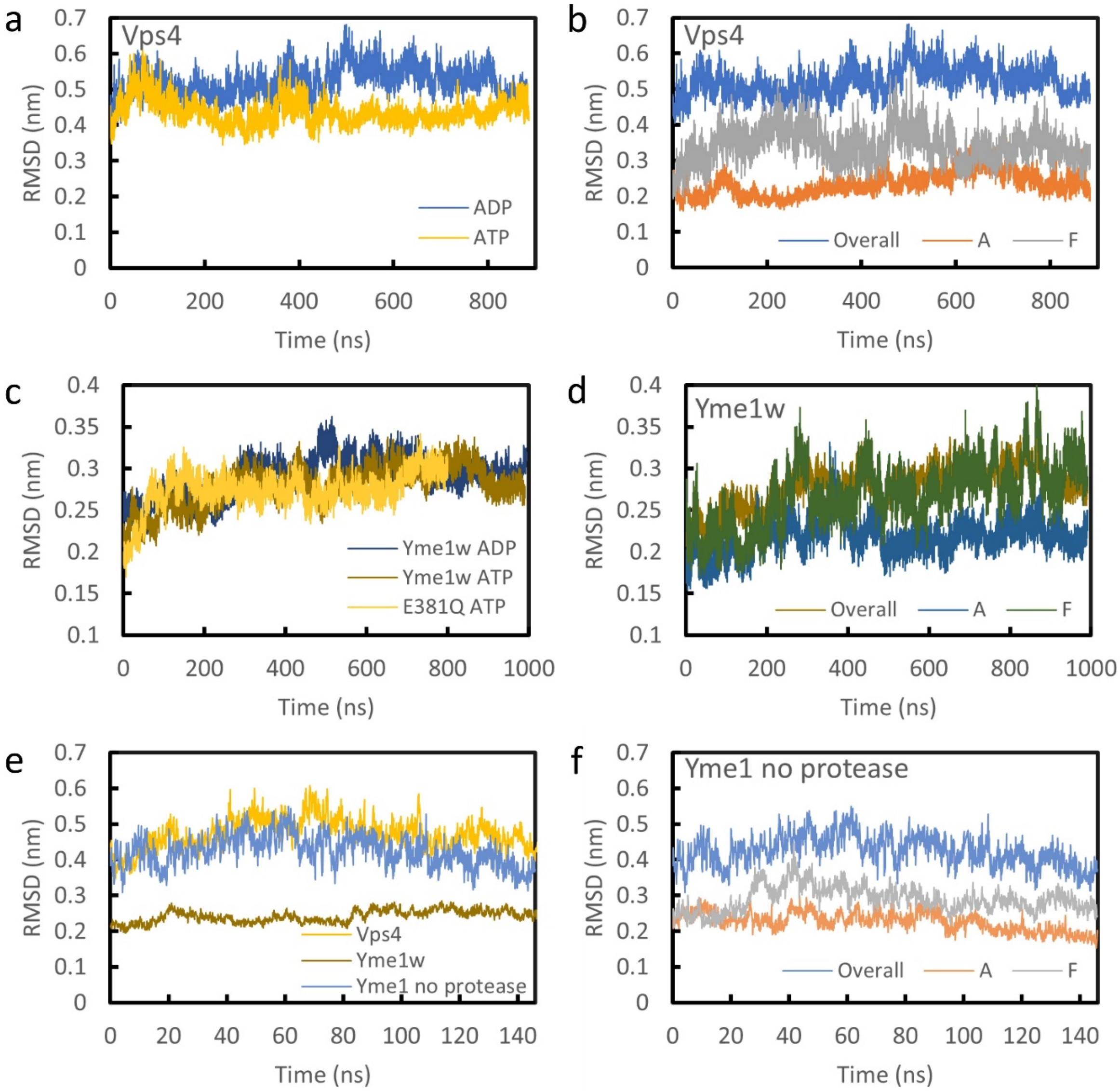


**Fig. S1.** Comparisons of Cα RMSD relative to cryo-EM structures for Vps4 and Yme1. **a)** Overall, full-hexamer RMSD of Vps4 with a polyL substrate peptide and ATP or ADP bound in subunit D. **b)** Overall and single-subunit RMSDs in Vps4 with a polyL substrate peptide and ADP-bound subunit D. **c)** Overall, full-hexamer RMSD for wild-type Yme1 (Yme1w) and the E381Q mutant binding either ATP or ADP in subunit D—each with a polyA substrate peptide. **d)** Overall and single-subunit RMSDs in wild-type Yme1 with a polyA substrate and ATP in subunit D. **e)** Overall RMSD for Vps4, wild-type Yme1, and truncated Yme1 without the protease domain. Vps4 had a polyL substrate and the Yme1 structures had a polyA substrate. All three structures had ATP in subunit D. **f)** Overall and single-subunit RMSD of protease-truncated Yme1 with a polyA substrate peptide and ATP in subunit D.

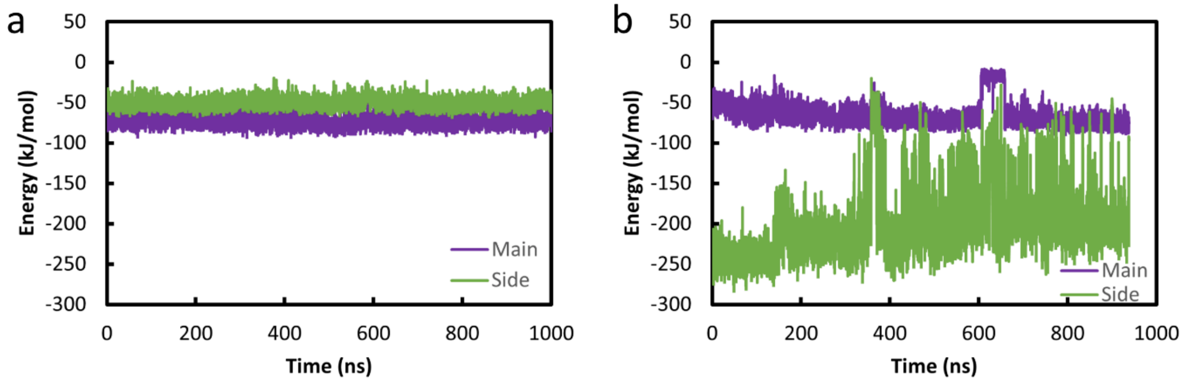


**Fig. S2.** Substrate peptide main- and sidechain contributions to interactions with subunit D pore loop 1 residues. Both plots are of Yme1 with subunit D binding ATP. **a)** PolyL substrate peptide. **b)** PolyVK substrate peptide.

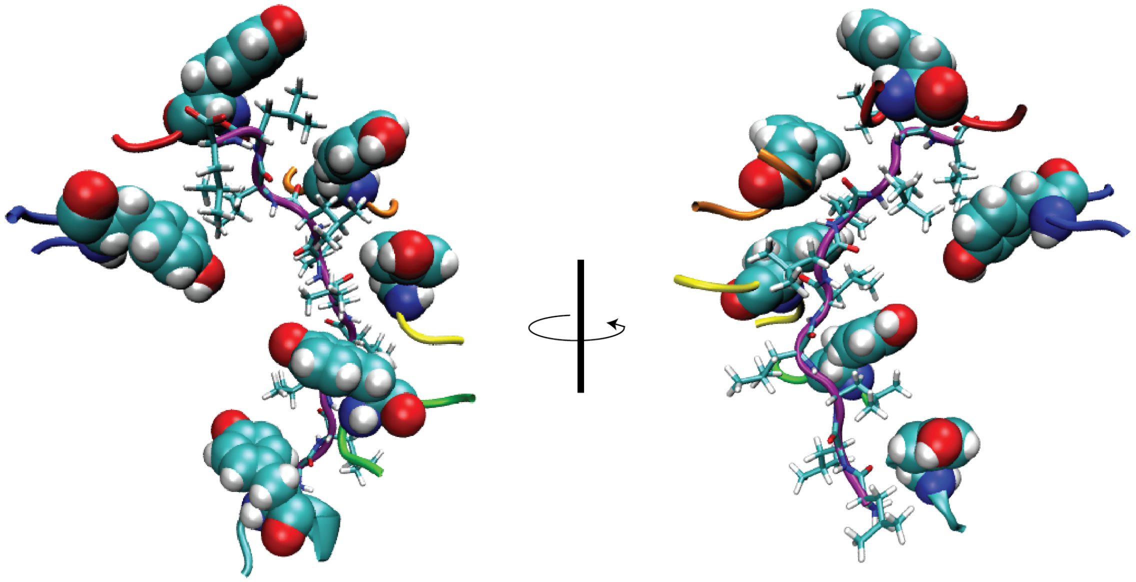

**Fig S3.** Tyr 354 of pore loop 1 intercalated between substrate side chains in YME1. Subunit pore loops are shown in their respective colors with Tyr shown with vdW radii. Fluctuating stacking and disengagement are shown for subunits E and F, respectively.

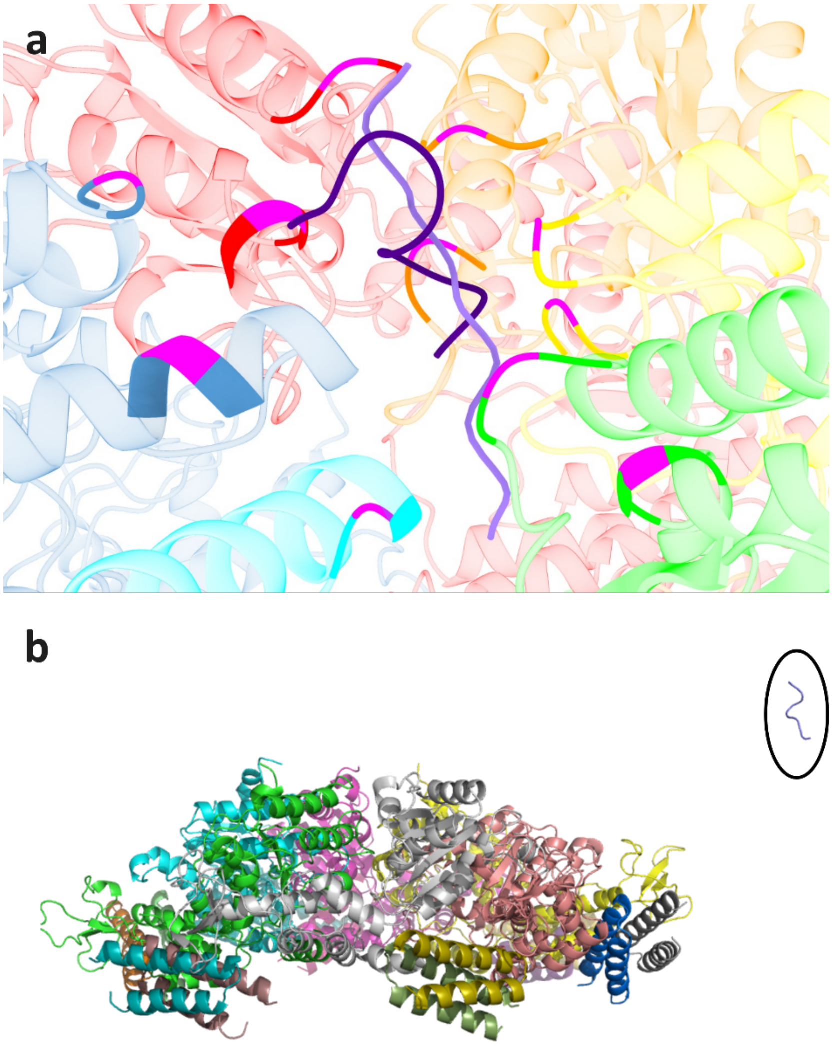


**Fig. S4.** Substrate peptides in reverse orientation. **a)** Overlay of Yme1 structure post simulation with cryo-EM-orientated (lighter purple) and reverse-oriented (darker purple) substrate peptides. Pore loop residues are colored magenta. Note that the reverse-oriented substrate has curled up like an alpha helix, indicating loss of interactions with the pore loops. **b)** A snapshot from the simulation of Vps4 with a reverse-orientation peptide circa 200ns. The substrate peptide has escaped from the pore and can be seen to the far right of the image (enclosed in a black circle for clarity).

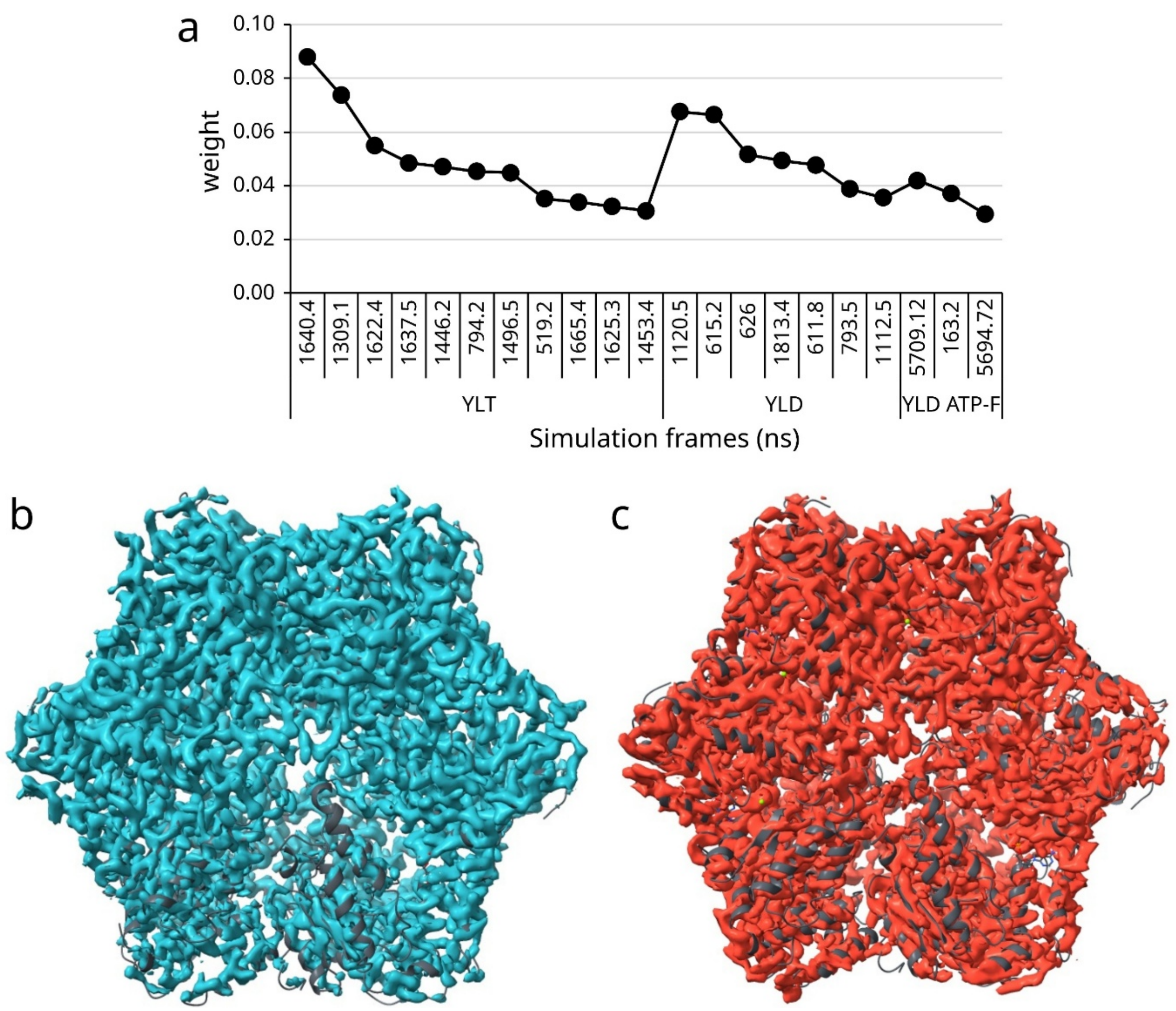


**Fig. S5.** cryoENsemble analysis results **a)** Weights of the 21 frames (numbered according to simulation time in ns) that best replicate the cryo-EM density based on cryoENsemble analysis. **b)** The cryo-EM density map for Yme1 with the cartoon representation of the cryo-EM structure superimposed in grey. Subunit F is in the lower-right portion of the map where the cartoon structure is most visible. This map was visualized at the recommended contour level of 0.08. **c)** The cryoENsemble-generated density map calculated from the weighted frames. The same cryo-EM cartoon as in b) is superimposed. The contour level of this map was set to enclose the same volume as the cryo-EM map in b).

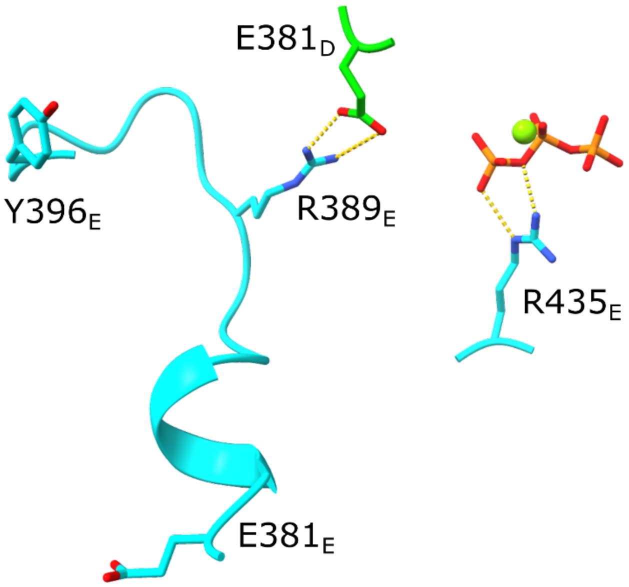


**Fig. S6.** The intersubunit loop connecting E381 to R389. This loop connects the hydrolysis-necessary Walker B E381 in one NBP to the previous subunit's pocket via R389. R389, in turn, coordinates the previous subunit's Walker B E381 in the pre-hydrolysis states. ATP phosphates, $Mg^{2+}$ ion, R435 arginine finger, and pore loop 2 Y396 are shown for reference.

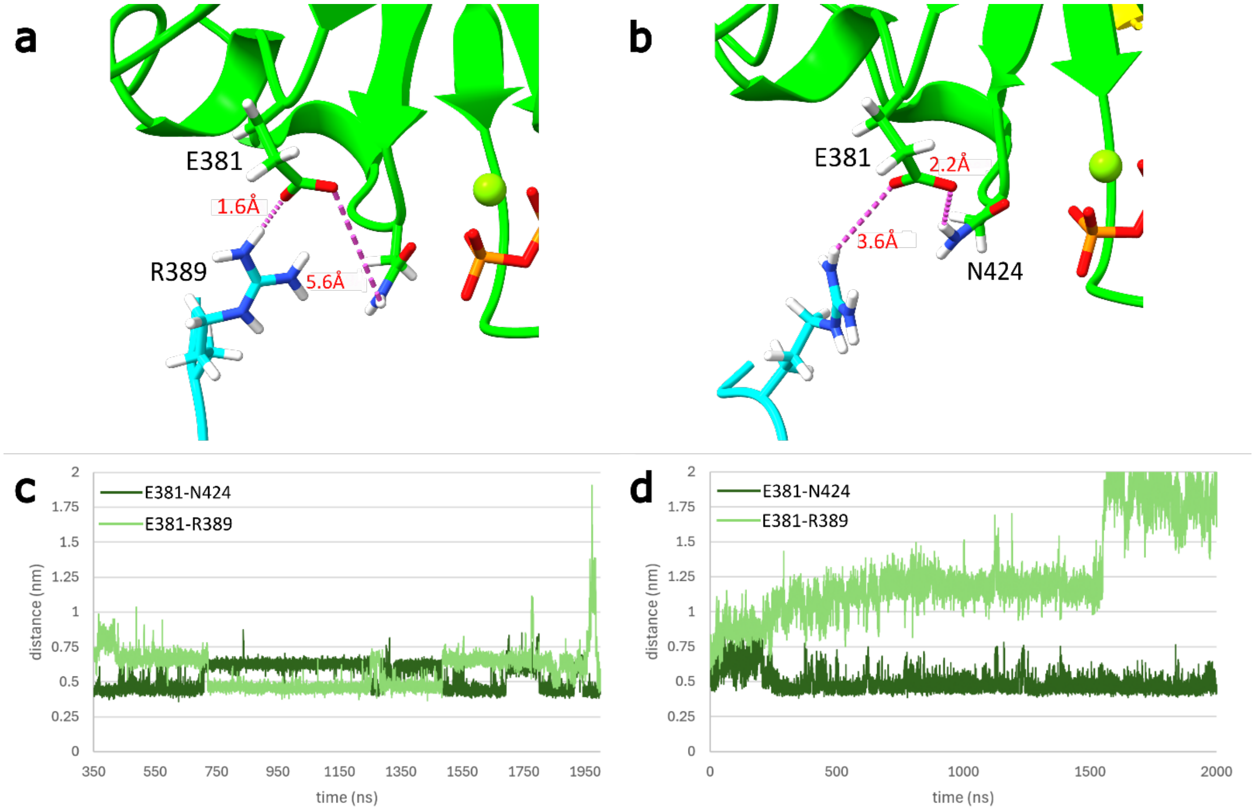


**Fig. S7.** Walker B interactions pre and post hydrolysis. **a)** Before hydrolysis Walker B E381 is salt bridged to R389 from the proximal subunit. This interaction is stable in subunits A-C. **b)** Once this salt bridge is disrupted, N424 stabilizes E381. **c)** At the C/D interface, the E381-R389 salt bridge is less stable, with E381 exchanging back and forth between R389 and N424 before hydrolysis (YLT), and **d)** stably engaging N424 post hydrolysis (YLD).

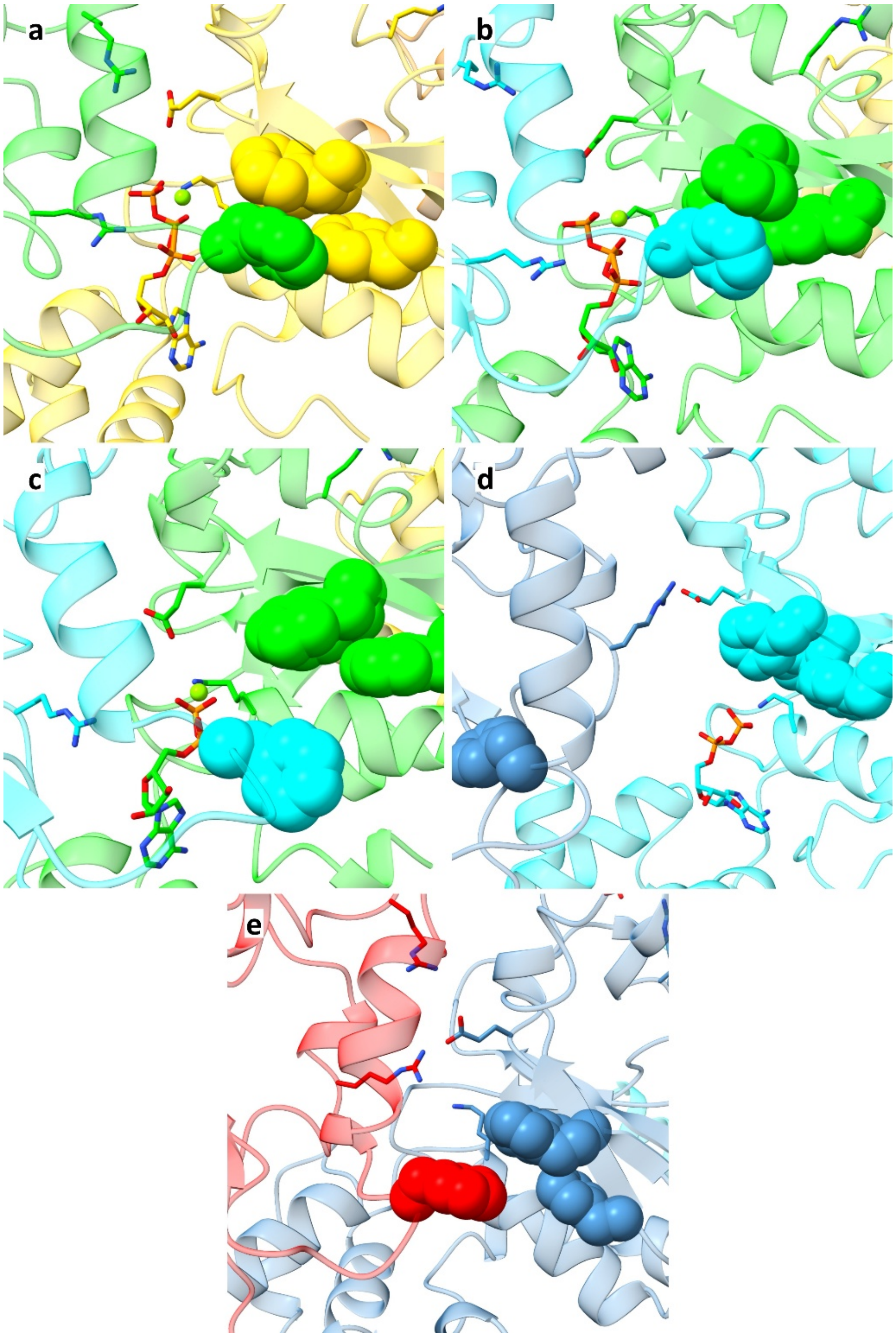


**Fig. S8.** ISS motif interactions throughout Yme1's mechanochemical cycle. The ISS motif residues and $Mg^{2+}$ ion are shown as spheres, with other NBP residues and the nucleotides shown as sticks. **a)** The first pre-hydrolysis state is shown in an upper subunit (subunit C). The ISS motif and other canonical intersubunit interactions are stably intact. **b)** The second pre-hydrolysis state in the hydrolysis-competent subunit D with ATP bound and in which the R389-E381 salt bridge is destabilized. ISS contacts remain largely stable, though some significant fluctuations are apparent. **c)** The first post-hydrolysis state shown in subunit D with ADP bound. The ISS motif fluctuates and separates as other intersubunit interactions—including R435-phosphate—are lost or significantly weakened. **d)** The second post-hydrolysis state is shown in the bottom subunit (subunit E). Here E381 has transiently salt bridged to the proximal R435 from F and the ISS motif cannot be reformed in this state as the residue from subunit F (F411) is stably incorporated into the C-terminus of helix α5. **e)** The resetting subunit F is shown in close contact with subunit A prior to ATP binding.

Now E381 is stably interacting with the proximal R435 from A while the ISS motif is available to reform contacts.

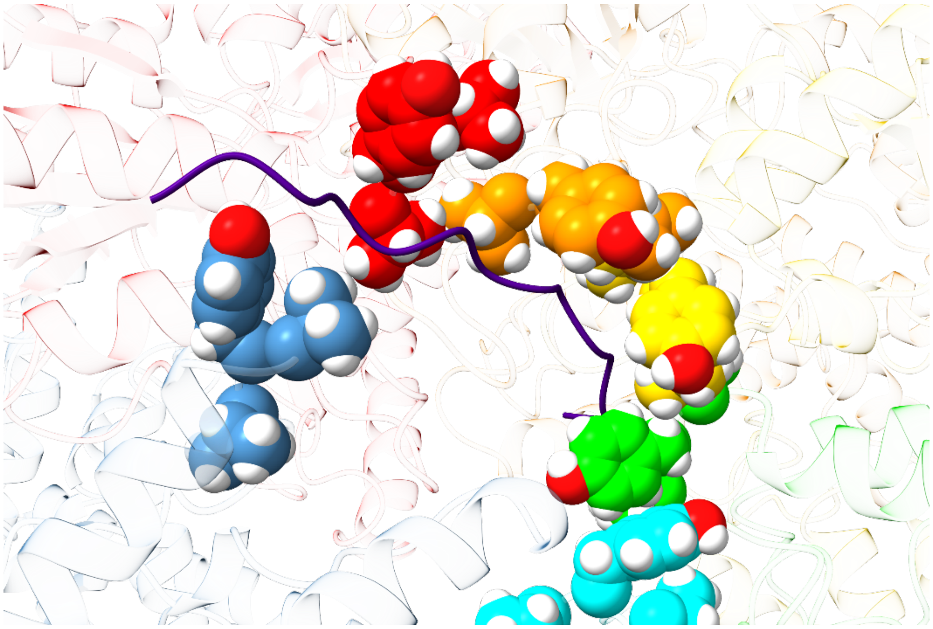

**Fig. S9.** View of the pore loop 1 residues V353-Y354-V355 as spheres and the bound substrate as a cartoon from YLD ATP-F simulation. Subunits A-D (red, orange, yellow, and green) are seen to interact in a regular manner with the substrate and each other. Subunit E (cyan) is dissociated from the substrate and the pore loop 1 spiral. Subunit F (blue) begins forming interactions with the substrate above subunit A.

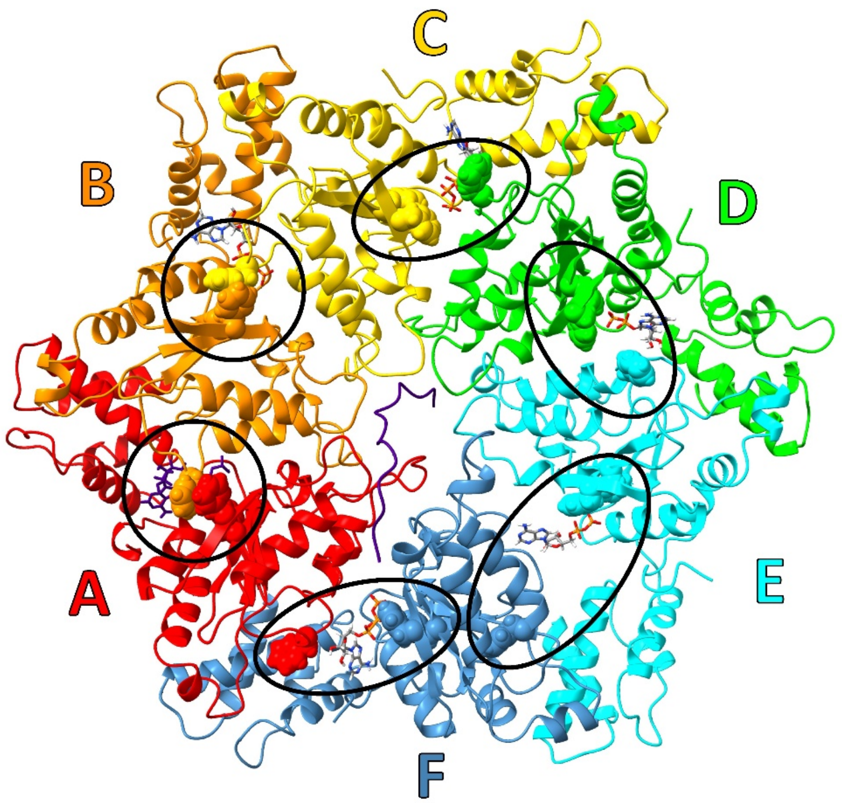


**Fig. S10.** An overhead view of Yme1 after 572.16 ns of simulation in the YLD ATP-F simulation. The ISS motif F411 and two of the opposing phenylalanines, F344 and F378, are shown as spheres and enclosed in black circles for clarity. Although formed for subunits A/B and B/C, the ISS interaction for C/D is lost, despite subunit C being in an ATP-bound state. This is reflective of additional strain in the ATPase and the potential for stochastic transitions.

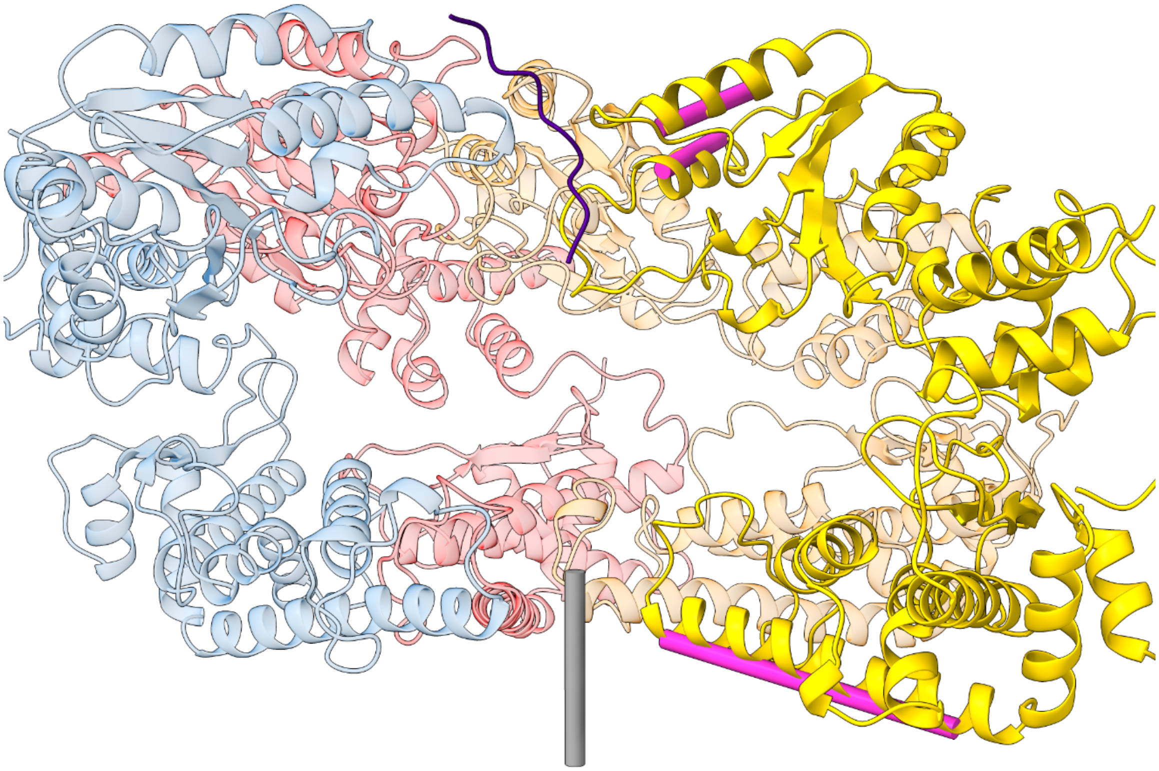

**Fig. S11.** Side view of the Yme1 hexamer, with two subunits removed for clarity. Axes used for ATPase-protease angle calculations are shown as rods. The magenta rods show the two ATPase helix vectors and the protease helix vector for one subunit. The gray rod represents the protease plane normal vector based on the six protease helix vectors.

### Tables

**Table S1.** Summary of systems simulated.

| ATPase | Substrate Peptide Sequence | Nucleotide State in Subunit D | Abbreviation | Run Time | GROMACS Version |
|---|---|---|---|---|---|
| Vps4 | cryo-EM[a] | ATP/ADP | | 200 ns | 2019.2 |
| | PolyL | ATP/ADP | VLT/VLD | ~880 ns | 2019.2 |
| | PolyVK | ATP/ADP | VVKT/VVKD | ~880 ns | 2019.2 |
| | Reverse | ATP | | 140 ns | 2019.4 |
| Vps4 | Total Simulation Time | | | ~4.1 µs | |
| Yme1 E381Q | cryo-EM (Poly-A) | ATP | | 800 ns | 2019.4 |
| Yme1 no protease | cryo-EM (Poly-A) | ATP | | ~150 ns | 2019.4 |
| Yme1 | cryo-EM (Poly-A) | ATP/ADP | YAT/YAD | 2 µs / 1.5 µs | 2019.4 |
| | PolyL | ATP/ADP | YLT/YLD | 2 µs | 2018.3 |
| | | ADP[b] | YLD ATP-F | 10 µs | N/A |
| | | ADP[b,c] | YLD Mg-E | 5 µs | N/A |
| | PolyVK | ATP | YVKT | ~940 ns | 2019.4 |
| | | ADP | YVKD | 2 µs | 2018.3 |
| | | ADP[b] | YVKD ATP-F | 10 µs | N/A |
| | Reverse | ATP | | 220 ns | 2019.4 |
| Yme1 | Total Simulation Time | | | ~36.6 µs | |

[a] An 8-residue ESCRT-III peptide with sequence ACE-ASP-GLU-ILE-VAL-ASN-LYS-VAL-LEU-$NH_2$

[b] ANTON2 production simulations with ATP docked in subunit F

[c] ANTON2 production simulation with ATP docked in subunit F and $Mg^{2+}$ added to subunit E

**Table S2.** Equilibration & production simulation steps. This table presents a breakdown of the multi-step equilibration process and production run with restraint force constants (FC), simulation time lengths, timestep, and ensemble.

| Stage | Step No. | Heavy Atom Restraint FC (kJ/mol/nm$^2$) | NBP Restraints[a] FC (kJ/mol/nm$^2$) | ANTON ATP-F and 14-Residue Substrate Restraints FC (kJ/mol/nm$^2$) | GROMACS Time (timestep) | ANTON Time (timestep) | Ensemble |
|---|---|---|---|---|---|---|---|
| Energy Minimization | 6.0 | 1000 | 0 | 500 | 5000 steps | 5000 steps | N/A |
| Freeze | N/A | Freeze | N/A | N/A | 100 ps (1 fs) | 100 ps (1 fs) | Berendsen thermostat |
| Equilibration | 6.1 w/ NBP[a] restraints | 1000 (500 for NBP restraints) | 500 | 500 | 0.5 ns (1 fs) | 0.5 ns (1 fs) | “ |
| | 6.2 | 800 | 0 | 500 | 0.5 ns (1 fs) | 0.5 ns (1 fs) | Berendsen NPT |
| | 6.3 | 600 | 0 | 500 | 0.5 ns (1 fs) | 0.5 ns (1 fs) | “ |
| | 6.4 | 400 | 0 | 500 | 1 ns (2 fs) | 1 ns (2 fs) | “ |
| | 6.5 | 200 | 0 | 500 | 10 ns (2 fs) | 1.5 ns (2 fs) | “ |
| | 6.6 | 100 (0 for subunit F) | 0 | 500 | 20 ns (2 fs) | 3 ns (2 fs) | “ |
| | 6.7 | 0 | 0 | 500 | 20 ns (2 fs) | 3 ns (2 fs) | “ |
| Production | 7 | 0 | 0 | 500 then 125[b,c] | Up to 2 µs (2 fs) | Up to 10 µs (2.4 fs) | GROMACS: V-rescale thermostat and Parrinello-Rahman barostat ANTON: Nose-Hoover thermostat and Martyna, Tuckerman, and Klein (MTK) barostat |

a) NBP restraints defined in Table S3
b) See Methods section
c) 14-residue substrate restraint only used during equilibration steps

**Table S3.** Individual restraint details. This table details the type and location of specific restraints used. Atoms are specified in the format "Residue:Atom", using the atom identifier from the PDB file.

| | Type | Restraint FC (kJ/mol) | Equilibrium distance | Atoms |
|---|---|---|---|---|
| NBP restraints | Harmonic | 500 | 0.2 nm | T328:OG1 – $Mg^{2+}$ |
| | | 500 | 0.195 nm | ATP:O2B - $Mg^{2+}$ |
| | | 500 | 0.195 nm | ATP:O2G - $Mg^{2+}$ |
| | | 500 | 0.195 nm | ATP:O3G - $Mg^{2+}$ |
| | Piecewise (GROMACS [bonds] type 10) | 500 at minimum distance | 0.45 nm minimum | E381:CD - $Mg^{2+}$ |
| Additional Anton ATP-F restraints | Harmonic | 500 | 0.624 nm | K327:NZ – ATP:PA |
| | | 500 | 0.381 nm | K327:NZ – ATP:PB |
| 14-residue substrate restraint | Piecewise (GROMACS [bonds] type 10) | 500 at minimum distance | 1.5 nm minimum | Residue 9:CA – Residue 14:CA |

**Movie legends**

**Movie S1 (separate file).** Animation of ATPase cycle. Composite of Movies S3-S9 showing multiple views of the Yme1 ATPase simultaneously.

**Movie S2 (separate file).** Animation of ATPase cycle. View of Yme1 pore loop 1 motifs.

**Movie S3 (separate file).** Animation of ATPase cycle. Top-down view of Yme1 hexamer.

**Movie S4 (separate file).** Animation of ATPase cycle. Top-down view of Yme1 subunit D NBP.

**Movie S5 (separate file).** Animation of ATPase cycle. Top-down view of Yme1 subunit E NBP.

**Movie S6 (separate file).** Animation of ATPase cycle. Top-down view of Yme1 subunit F NBP.

**Movie S7 (separate file).** Animation of ATPase cycle. Side view of Yme1 subunit D NBP.

**Movie S8 (separate file).** Animation of ATPase cycle. Side view of Yme1 subunit E NBP.

**Movie S9 (separate file).** Animation of ATPase cycle. Side view of Yme1 subunit F NBP.

### SI References


1. H. M. Berman, J. Westbrook, Z. Feng, G. Gilliland, T. N. Bhat, H. Weissig, I. N. Shindyalov, P. E. Bourne, The Protein Data Bank. *Nucleic Acids Res.* **28**, 235–242 (2000).
2. B. Webb, A. Sali, Comparative Protein Structure Modeling Using MODELLER. *Curr. Protoc. Bioinformatics* **54**, 5.6.1-5.6.37 (2016).
3. A. Fiser, R. Kinh, G. Do, A. S. ˇ Ali, Modeling of loops in protein structures. *Protein Science* **9**, 1753–1773 (2000).
4. M. H. M. Olsson, C. R. SØndergaard, M. Rostkowski, J. H. Jensen, PROPKA3: Consistent treatment of internal and surface residues in empirical p K a predictions. *J. Chem. Theory Comput.* **7**, 525–537 (2011).
5. K. Jonas, J. Liu, P. Chien, M. T. Laub, Proteotoxic Stress Induces a Cell-Cycle Arrest by Stimulating Lon to Degrade the Replication Initiator DnaA. *Cell* **154**, 623–636 (2013).
6. A. J. Rampello, S. E. Glynn, Identification of a Degradation Signal Sequence within Substrates of the Mitochondrial i-AAA Protease. *J. Mol. Biol.* **429**, 873–885 (2017).
7. H. Shi, A. J. Rampello, S. E. Glynn, Engineered AAA+ proteases reveal principles of proteolysis at the mitochondrial inner membrane. *Nature Communications 2016 7:1* **7**, 1–12 (2016).
8. M. J. Abraham, T. Murtola, R. Schulz, S. Páll, J. C. Smith, B. Hess, E. Lindah, GROMACS: High performance molecular simulations through multi-level parallelism from laptops to supercomputers. *SoftwareX* **1–2**, 19–25 (2015).
9. H. J. C. Berendsen, D. van der Spoel, R. van Drunen, GROMACS: A message-passing parallel molecular dynamics implementation. *Comput. Phys. Commun.* **91**, 43–56 (1995).
10. D. E. Shaw, J. P. Grossman, J. A. Bank, B. Batson, J. A. Butts, J. C. Chao, M. M. Deneroff, R. O. Dror, A. Even, C. H. Fenton, *et al.*, Anton 2: Raising the Bar for Performance and Programmability in a Special-Purpose Molecular Dynamics Supercomputer. *International Conference for High Performance Computing, Networking, Storage and Analysis, SC* **2015-January**, 41–53 (2014).
11. J. Huang, S. Rauscher, G. Nawrocki, T. Ran, M. Feig, B. L. De Groot, H. Grubmüller, A. D. MacKerell, CHARMM36m: an improved force field for folded and intrinsically disordered proteins. *Nature Methods 2016 14:1* **14**, 71–73 (2016).
12. J. Huang, A. D. Mackerell, CHARMM36 all-atom additive protein force field: Validation based on comparison to NMR data. *J. Comput. Chem.* **34**, 2135–2145 (2013).
13. W. L. Jorgensen, J. Chandrasekhar, J. D. Madura, R. W. Impey, M. L. Klein, Comparison of simple potential functions for simulating liquid water. *J. Chem. Phys.* **79**, 926–935 (1983).
14. H. J. C. Berendsen, J. P. M. Postma, W. F. Van Gunsteren, A. Dinola, J. R. Haak, Molecular dynamics with coupling to an external bath. *J. Chem. Phys.* **81**, 3684–3690 (1984).
15. B. Hess, P-LINCS:  A Parallel Linear Constraint Solver for Molecular Simulation. *J. Chem. Theory Comput.* **4**, 116–122 (2007).
16. M. Parrinello, A. Rahman, Polymorphic transitions in single crystals: A new molecular dynamics method. *J. Appl. Phys.* **52**, 7182–7190 (1981).
17. S. Nosé, A molecular dynamics method for simulations in the canonical ensemble. *Mol. Phys.* **52**, 255–268 (1984).
18. W. G. Hoover, Canonical dynamics: Equilibrium phase-space distributions. *Phys. Rev. A (Coll. Park).* **31**, 1695 (1985).
19. G. J. Martyna, D. J. Tobias, M. L. Klein, Constant pressure molecular dynamics algorithms. *J. Chem. Phys.* **101**, 4177–4189 (1994).
20. W. Humphrey, A. Dalke, K. Schulten, VMD: Visual molecular dynamics. *J. Mol. Graph.* **14**, 33–38 (1996).
21. E. C. Meng, T. D. Goddard, E. F. Pettersen, G. S. Couch, Z. J. Pearson, J. H. Morris, T. E. Ferrin, UCSF ChimeraX: Tools for structure building and analysis. *Protein Science* **32**, e4792 (2023).
22. E. F. Pettersen, T. D. Goddard, C. C. Huang, E. C. Meng, G. S. Couch, T. I. Croll, J. H. Morris, T. E. Ferrin, UCSF ChimeraX: Structure visualization for researchers, educators, and developers. *Protein Science* **30**, 70–82 (2021).
23. N. Michaud-Agrawal, E. J. Denning, T. B. Woolf, O. Beckstein, MDAnalysis: A toolkit for the analysis of molecular dynamics simulations. *J. Comput. Chem.* **32**, 2319–2327 (2011).

24. R. J. Gowers, M. Linke, J. Barnoud, T. J. E. Reddy, M. N. Melo, S. L. Seyler, J. Domański, D. L. Dotson, S. Buchoux, I. M. Kenney, *et al.*, MDAnalysis: A Python Package for the Rapid Analysis of Molecular Dynamics Simulations. *scipy* 98–105 (2016). https://doi.org/10.25080/MAJORA-629E541A-00E.
25. T. Włodarski, J. O. Streit, A. Mitropoulou, L. D. Cabrita, M. Vendruscolo, J. Christodoulou, Bayesian reweighting of biomolecular structural ensembles using heterogeneous cryo-EM maps with the cryoENsemble method. *Sci. Rep.* **14**, 18149 (2024).
26. C. Puchades, A. J. Rampello, M. Shin, C. J. Giuliano, R. L. Wiseman, S. E. Glynn, G. C. Lander, Structure of the mitochondrial inner membrane AAA+ protease YME1 gives insight into substrate processing. *Science.* **358**, eaao0464 (2017).
